# Tuberculostearic acid (TSA)-containing phosphatidylinositols as reliable marker to determine *Mycobacterium tuberculosis* bacterial burden

**DOI:** 10.1101/2021.02.04.429149

**Authors:** Julius Brandenburg, Jan Heyckendorf, Franziska Waldow, Nicole Zehethofer, Lara Linnemann, Nicolas Gisch, Hande Karaköse, Maja Reimann, Katharina Kranzer, Barbara Kalsdorf, Patricia Sanchez-Carballo, Michael Weinkauf, Verena Scholz, Sven Malm, Susanne Homolka, Karoline I. Gaede, Christian Herzmann, Ulrich E. Schaible, Christoph Hölscher, Norbert Reiling, Dominik Schwudke

## Abstract

It is estimated that approximately one-fourth of the world's population is infected with strains of the *Mycobacterium tuberculosis* complex (MTBC), the causative agents of tuberculosis (TB). In this study, we present rationally developed molecular markers for bacterial burden, which are derived from mycobacterial phospholipids. Using lipidomic approaches, we show that tuberculostearic acid (TSA)-containing phosphatidylinositols (PI) are present in all clinically relevant MTBC lineages investigated. For the major abundant lipid PI 16:0_19:0 (TSA), a detection limit equivalent to 10^2^ colony forming units (CFU) was determined for bacterial cultures and approximately 10^3^ for cell culture systems. We further developed a mass spectrometry based targeted lipid assay, which – in contrast to bacterial quantification on solid medium – can be performed within several hours including sample preparation. Translation of this indirect and culture-free detection approach allowed the determination of pathogen loads in infected murine macrophages, human neutrophils and murine lung tissue. We show that marker lipids inferred from the mycobacterial PIs are increased in peripheral blood mononuclear cells (PBMCs) of TB patients beyond the lipid metabolic background in comparison to healthy controls. In a small cohort of drug-susceptible TB patients elevated levels of these marker molecules were detected at therapy start and declined following successful anti-tuberculosis treatment. The concentration of TSA-containing PIs can be used as correlate for reliable and rapid quantification of *Mycobacterium tuberculosis (Mtb)* burden in experimental *in vitro* model systems and may also provide a clinically relevant tool for monitoring TB therapy.

**One Sentence Summary:** Tuberculostearic acid containing phosphatidylinositols represent a novel, fast to measure, reliable correlate of *Mycobacterium tuberculosis* bacterial burden in experimental model systems, which makes a future clinical application conceivable.

## Introduction

Globally, tuberculosis (TB) remains a leading cause of morbidity and mortality with more than 10 million active cases annually (*1*). The causative agent, *Mycobacterium tuberculosis (Mtb),* is responsible for approximately 1.5 million deaths per year. Emerging drug-resistance such as multidrug-resistant (MDR – defined by resistance against rifampicin and isoniazid) TB cases is alarming since treatment fails in a high portion of such patients (*2, 3*). From this perspective, biomarkers that reflect therapy responses are highly desirable to rapidly adjust a treatment to the individual patient's medical need (*4*). In this perspective, lipid markers of mycobacterial origin may represent ideal candidates to assess bacterial burden in general.

The cell envelope of *Mycobacterium tuberculosis* complex (MTBC) bacteria is complex in nature and consists of a plasma membrane (PM) as the innermost layer that is covered by the peptidoglycan (PGN)-arabinogalactan (AG) complex. The AG is further esterified by long-chain mycolic acids (MA). Into these MA various complex lipids and lipoglycans such as sulfolipids (SL), trehalose dimycolates (TDM), phenolic glycolipids (PGL), lipoarabinomanan (LAM) and phthiocerol dimycocerosates (PDIM) are non-covalently intercalated, thus forming a strong outer envelope, the so-called mycomembrane (MM) (*5, 6*). The mycobacterial PM mainly consists of phospholipids and is similar to those seen in other microorganisms (*7*). Their main components are phosphatidyl-*myo*-inositol mannosides (PIM), phosphatidylinositol (PI), phosphatidylethanolamine (PE), phosphatidylglycerol (PG), and cardiolipin (CL). The major fatty acid constituents of the PM are palmitic (16:0), octadecenoic (18:1) and 10-methyloctadecanoic (tuberculostearic acid – TSA, (19:0)). Mycobacterial lipids represent a pool of potential biomarkers to trace the status of *Mtb*-infections due to their unique structural features not found in eukaryotic lipidomes (*8*). Due to high diversity and complexity of these lipids, approaches to measure and profile them are challenging (*9*). Various mass spectrometry (MS)-based lipidomics approaches have been developed during the last 15 years to foster the analysis of lipids further satisfying the rising interest on this class of biomolecules (*10, 11*). MAs have been of special interest, however, since standards are ill-defined, the development of MA-based diagnostic markers remains difficult (*12, 13*). The complex lipids of *Mtb* are comparably large molecules and mostly comprise a neutral net charge, which require specific extraction procedures as well as ionization and detection conditions (*8*). In comparison, for phospholipids numerous and very advanced analytic procedures have been established. This is reflected in a recent study, which showed that different PIM and PI species enable MALDI mass spectrometry imaging (MALDI-MSI) of intact *Mtb* in rabbit-lung granulomas (*14*).

In the current study, we identified saturated mycobacterial PIs containing TSA as a novel and reliable marker of bacterial loads in relevant TB infection models *in vitro* and *in vivo.* We further observed that specific PIs were significantly enriched in PBMCs from active TB patients, which may open a future perspective for its use in a clinical setting.

## Results

### Phosphatidylinositols containing TSA represent reliable lipid markers for mycobacterial load

Our search for lipid-based markers of MTBC strains was focused on amphiphilic molecules that should be extractable with established methods, easy to quantify and present in the pathogen with high abundance. Therefore, we profiled the content of the methanolic fraction by liquid chromatography-mass spectrometry (LC-MS) to perform a targeted analysis of PM lipids (fig. S1). PL profiles of 18 MTBC strains covering five lineages, both ancient and modern ones including three clinical Beijing isolates, exhibited a high similarity (Table 1, Fig. 1). We identified as main molecular species, PE and PI comprising a diacylglycerol (DAG) backbone built from FA 16:0 and FA 19:0 (TSA) for all investigated MTBC strains (Fig. 1A, B). Presence of TSA as the sole fatty acid isomer comprising a 19:0 acyl chain was proven by gas-chromatography coupled with mass spectrometry (GC-MS) analysis of the fatty acid methyl ester (FAME) profiles generated from the methanolic fraction (data not shown), being in line with literature (*15*). PE 35:0 (16:0_19:0 (TSA)) is the most abundant molecular species of its class with an average of 31.9 mol%, but its abundance varied to a larger extent between the investigated isolates. PI 35:0 (16:0_19:0 (TSA)) was quantified as major abundant molecular species of its class in all strains with 70.7 mol% on average (Fig. 1B, C, fig. S3). It is noteworthy that for the PG class PG 35:0 (16:0_19:0 (TSA)) was only a minor component, while PG 34:1 was the most abundant species (fig. S2). TSA was further detected in combination with FA 14:0, FA 15:0, FA 16:1, FA 17:0, FA 17:1 and FA 18:0 in PE and PI, as well as LPE 19:0 (lysophosphatidylethanolamine) and LPI 19:0 (lysophosphatidylinositol) in these strains (data file S1). PI 16:0_19:0 (TSA) has the analytical advantage that additional fragments of the PI headgroup can be detected by tandem mass spectrometry (MS^2^) with high sensitivity and specificity, which is not the case for PE 16:0_19:0 (fig. S4). Thus, PI 16:0_19:0 (TSA) was further studied in detail.

**Table 1.**
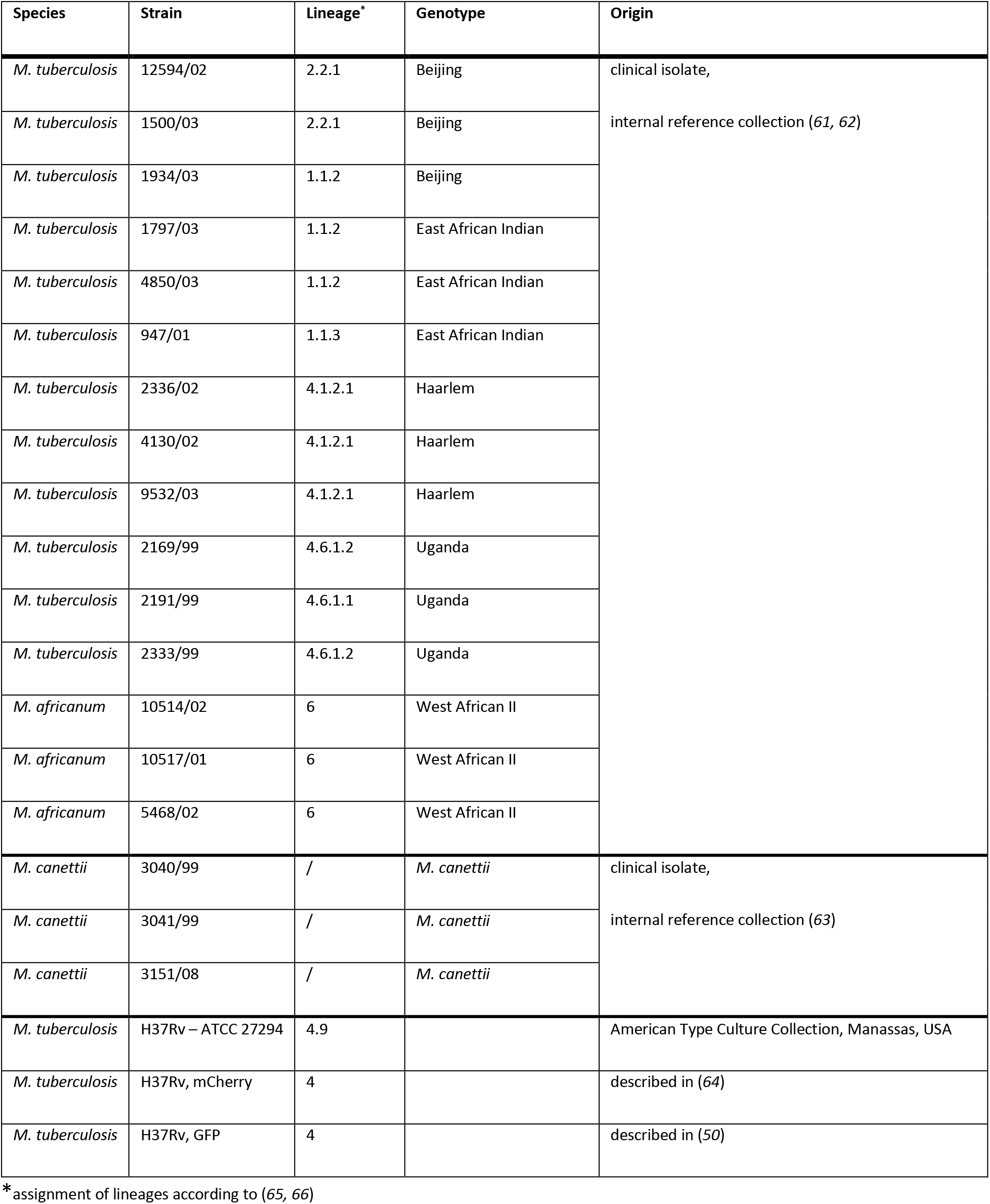
Phylogenetic classification of all investigated MTBC strains.

**Fig. 1.**
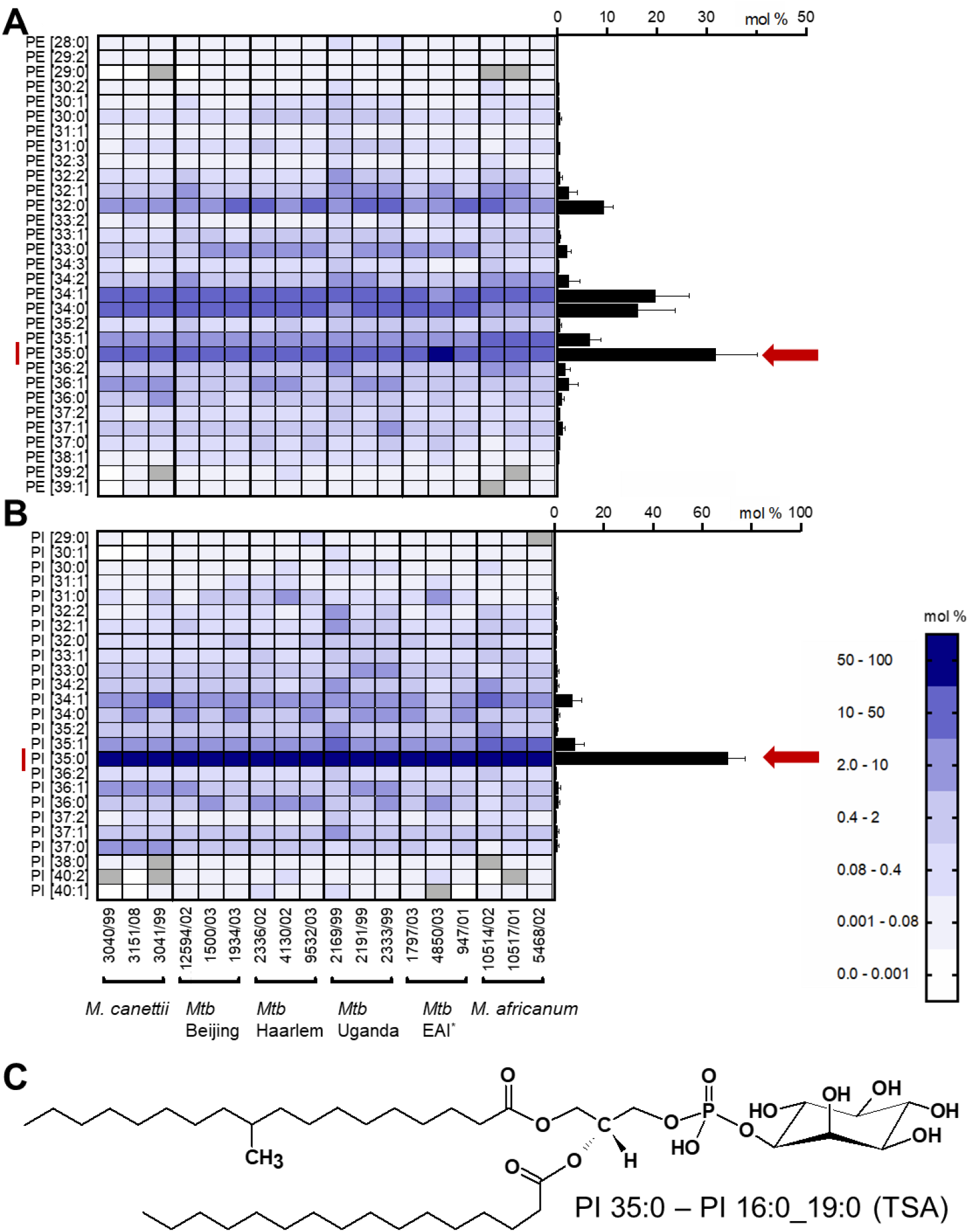
PI 35:0 – 16:0_19:0 (TSA) is the abundant phosphoglycerolipid of MTBC clinical isolates. Lipid profiles of **A)** phosphatidylethanolamine (PE) and **B)** phosphatidylinositol (PI) were determined using high resolution LC-MS from extracts of clinical MTBC isolates. Profiles of clinical isolates were determined from at least two independent cultivations (separate profiles for each strain are shown in fig. S2). On the left a heat map representation for individual strains is provided. The average lipid profiles for all strains are represented in the right as bar graph (Error bars represent one SD). **C)** Structure of the *sn*-1 isomer of PI 16:0_19:0 (TSA) used as marker for mycobacterial loads and CFU. *East African Indian strains – EAI.

Subsequently, we evaluated whether the PI 16:0_19:0 (TSA) concentration could be used to estimate colony forming units (CFUs) using dilutions of three independent cultures of *Mtb* H37Rv mCherry. Lipid analysis of bacterial pellets was completed within 24 hours including inactivation of *Mtb.* We observed a linear correlation between the amount of PI 16:0_19:0 (TSA) and CFU (Fig. 2A). Although 100 mycobacteria were still detectable, the lower limit of quantitation for the CFU determination by the PI 16:0_19:0 (TSA) assay was approximately 1,000 bacteria. This result indicates that targeted quantitation of PI 16:0_19:0 (TSA) is a fast and sensitive way to determine CFU of *Mtb*. From this data, we can estimate that a single mycobacterium contains approximately 45 amol of PI 16:0_19:0 (TSA). This finding is in good agreement with the detection limit of the lipidomics platform for PI molecular species of approximately 1 fmol. The feasibility of this approach was further evaluated in an independent bacteriological test system for sputum analysis. Here, we demonstrated that a mycobacterial load of 2,000 CFU could be detected in both, artificial sputum and sputum supplemented with a background flora composed of fungal and bacterial species (Fig. 2B).

**Fig. 2.**
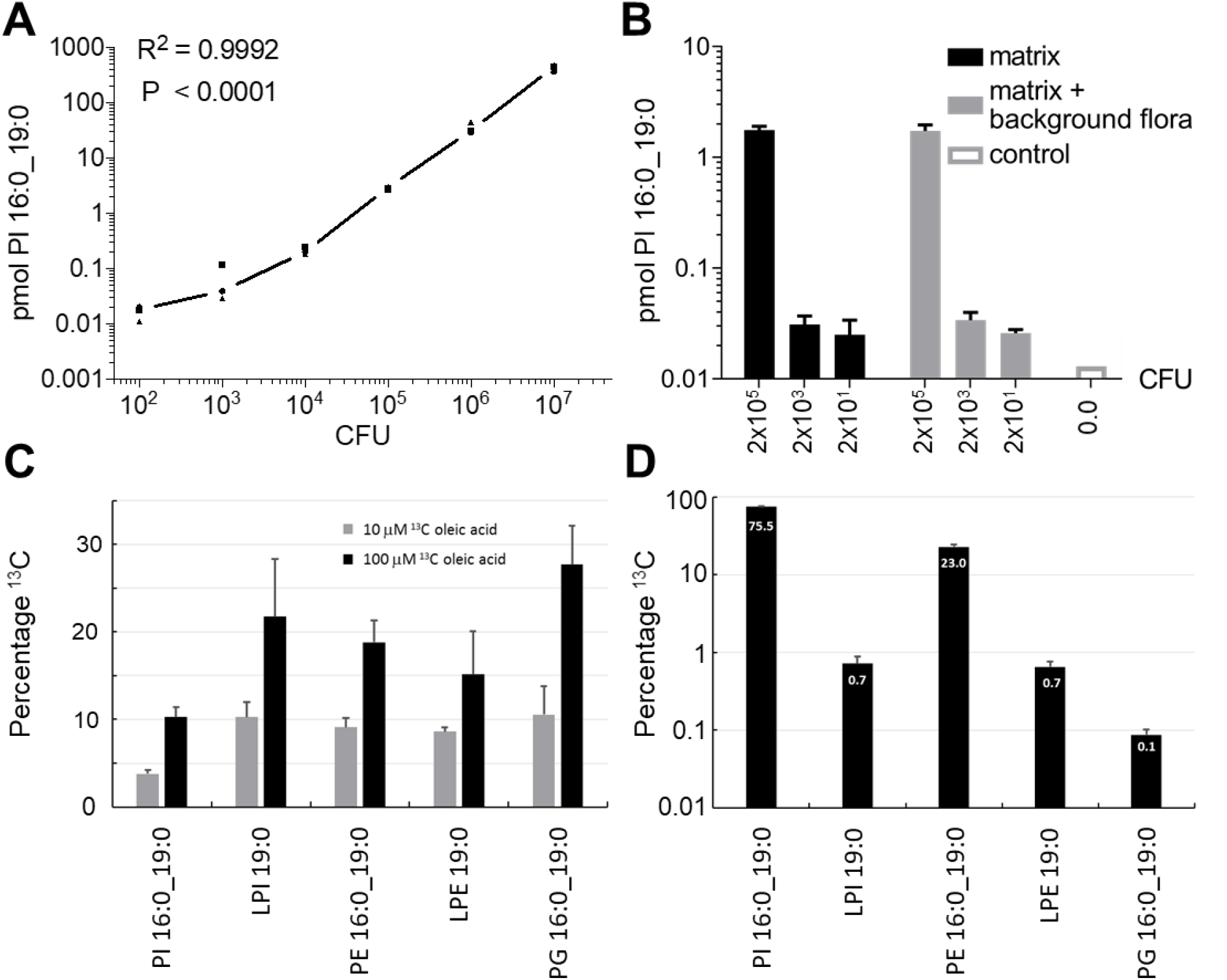
Detection of *Mtb* in culture using PI 16:0_19:0 and metabolic labelling of TSA. **A)** Correlation between the amount of PI 16:0_19:0 (TSA) and CFU (n=3) of *Mtb* H37Rv (mCherry). **B)** Detection of *Mtb* in artificial sputum in presence of bacterial and fungal flora using PI 16:0_19:0 (TSA). **C)** Labelling efficiency of *Mtb* using ^13^C-labeled oleic acid (FA 18:1, OA) which is metabolized to FA 19:0 (TSA) by the pathogen and **D)** relative abundance of main FA 19:0 (TSA)-containing membrane lipids.

### Tracer analysis using ^13^C-labelled oleic acid and its transformation to TSA enables monitoring of Mtb growth

The biosynthesis of the PM is crucial for *Mtb* growth. Therefore, a tracer analysis of its biosynthesis will provide a well-defined estimate for mycobacterial replication. As substrate molecule oleic acid (OA; FA 18:1) was chosen, a highly abundant building block of membrane lipids and neutral lipids in mammalian cells. When the growth medium was supplemented with ^13^C18-labelled OA (^13^C18 OA), we observed its transformation to TSA by addition of a methyl-group comprising one ^12^C atom (fig. S5–8). Such labelled TSA served as building block for the membrane lipids, PI 16:0_19:0 (TSA), PE 16:0_19:0 (TSA) and PG 16:0_19:0 (TSA) as well as LPI 19:0 (TSA) and LPE 19:0 (TSA) (Fig. 2C, fig. S5–8). After seven days of incubation, we observed dose-dependent labelling efficiencies for PI 16:0_19:0 (TSA) of 3.8% with a supplement of 10 μM ^13^C18 OA, which was further increased to 10.3% for 100 μM for the tracer molecules. The labelling efficiency for LPI 19:0 (TSA), PE 16:0_19:0 (TSA), LPE 19:0 (TSA) and PG 16:0_19:0 (TSA) was approximately doubled for both conditions when compared to PI 16:0_19:0 (TSA), but in terms of total amounts PI 16:0_19:0 (TSA) was the major labelled species (Fig. 2D). An even higher labelling efficiency for PI 16:0_19:0 was recently observed when *Mtb* was cultured together with macrophages previously labelled with ^13^C18 OA (preprint:(*16*)).

### Quantitation of PI 16:0_19:0 enables to monitor the Mtb infection status in cell culture systems

Next, we used MS/MS to perform a targeted analysis of PI 16:0_19:0 (TSA) to monitor all isomers in the mass range of 851.5 +/-0.5 Da (fig. S4) in order to address the infection status and the mycobacterial load in cell culture systems. For 36 independent cultures of murine bone marrow derived macrophages (BMDM), the infection status was correctly determined with only one false negative test (Fig. 3A). *Mtb* infection status was determined using two mass spectrometric signals, the precursor ion at *m/z* 851.5658 and the FA 19:0 (TSA) fragment ion 297.2798 (fig. S5A). The applicability of PI 16:0_19:0 as a marker of mycobacterial load was further analyzed in infection experiments with human neutrophils. Neutrophils from five healthy donors were infected with a multiplicity of infection (MOI) of 3 and incubated for 6 hours. After this short incubation time phagocytosis is completed but intracellular pathogens have not yet multiplied nor were the mycobacteria killed by the neutrophils (*17*). This approach enabled us to evaluate if a basal concentration of PI 16:0_19:0 (TSA) was present in human neutrophils. All control cultures were identified as negative (n = 11), and all cultures infected with *Mtb* as positive (ion at *m/z* 851.5658 in MS^1^ and fragment ion 297.2798 in MS^2^ detectable; n = 9) (Fig. 3B). The experimental variation for this standardized infection model was further compared by relative standard deviation (RSD). Lipid quantitation had a much better precision with RSD of 0.20 than CFU determination with RSD of 0.70 (Fig. 3B). These results indicated that PI 16:0_19:0 (TSA) is specific enough to detect and quantify *Mtb* load in microbiological experiments and murine and human cell culture models.

**Fig. 3.**
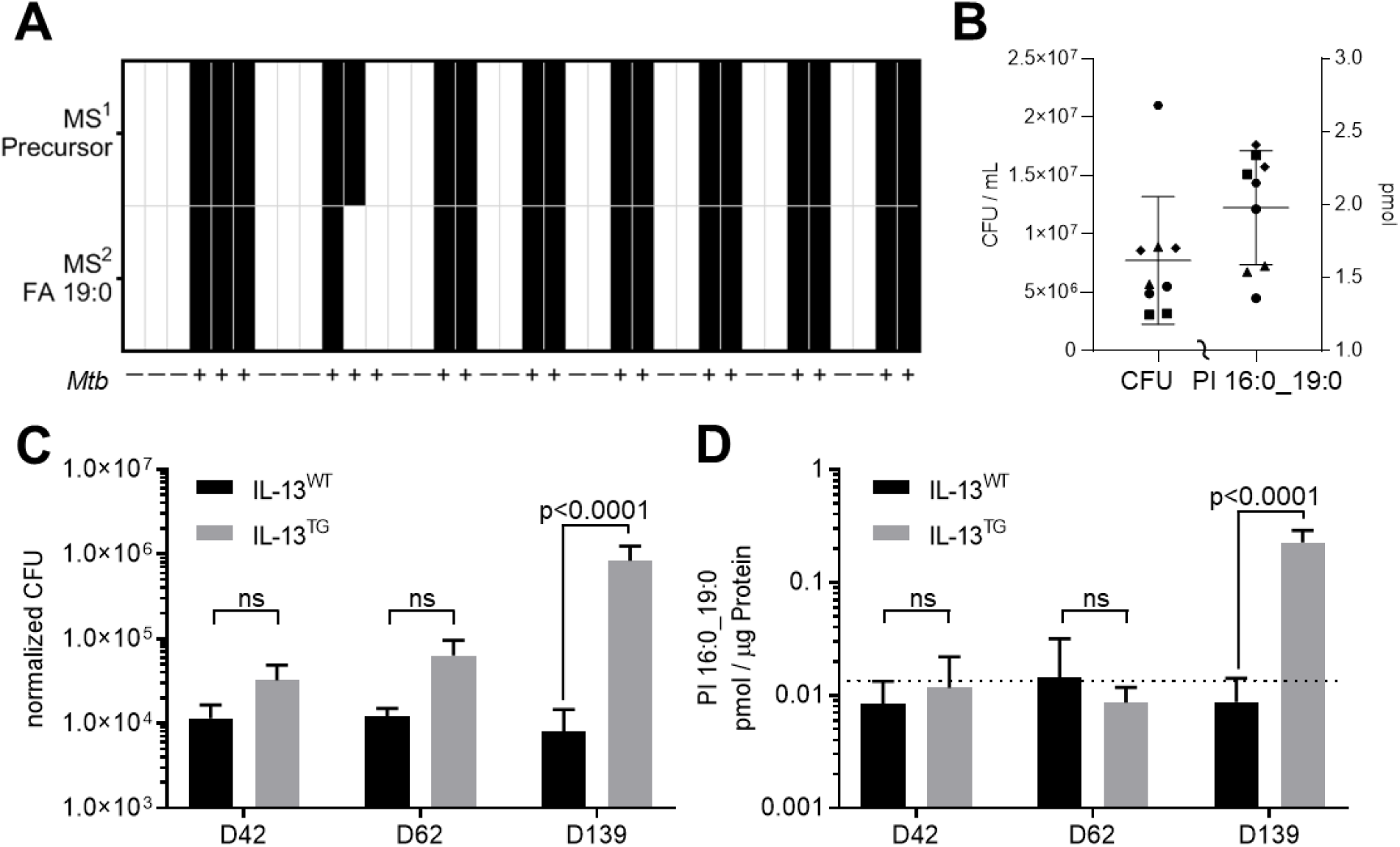
Application of PI 16:0_19:0 as correlate of *Mtb* load in model systems. **A)** Detection of *Mtb* in macrophage cultures. MS detection level either on basis of accurate mass determination in MS^1^ (*m/z* 851.5655) or MS^2^ (FA 19:0, *m/z* 851.5 -> 297.2799 – fig. S4A) **B)** Validation of *Mtb* infection status for neutrophils isolated from 5 healthy donors. At 6 hpi CFU (left axis) and amount of PI 16:0_19:0 (right axis) were determined. Data from different donors are indicated by distinctive symbols with double determination for each individual. In non-infected cultures PI 16:0_19:0 was below the limit of detection. **C)** Pulmonary mycobacterial burden in IL-13^WT^ and IL-13^TG^ mice upon *Mtb* aerosol infection by CFU analysis at indicated time points and **D)** respective concentrations of PI 16:0_19:0 determined from the same lung tissue homogenates. Dotted line indicated basal concentration in uninfected mice (1.75×10^-2^ pmol/μg; n=11, data file S2).

### Monitoring CFU in Mtb infected mice by quantitation of PI 16:0_19:0

Many studies on fundamental processes during *Mtb* infection are performed in experimental mouse models. Of particular interest are models, which reflect TB disease in humans (*18, 19*). This holds also true for IL-13^TG^ animals, in which *Mtb* infection – in contrast to IL-13^WT^ (C57Bl/6) mice – leads to unrestricted bacterial growth and subsequent development of centrally necrotizing granulomas, which strongly resemble the morphology of lesions in TB patients (*20, 21*). In the present study, we compared the growth of *Mtb* in IL-13^TG^ and IL-13^WT^ mice and compared CFU development and the presence of PI 16:0_19:0 at day 42, 62 and 139 post infection (p.i.). We observed a significant increase in bacterial numbers in lungs of IL-13^TG^ when compared to IL-13^WT^ mice 139 days after infection (Fig. 3C). In *Mtb* infected IL-13^WT^ control mice CFU levels measured at d42, d62 and d139 stayed at approximately 1.0×10^4^ bacteria per extracted tissue sample (4.7×10^6^ CFU / g tissue, data file S2) during *Mtb* infection, which corroborates previous studies that CFU levels in the lung of C57Bl/6 mice remain rather stable during chronic experimental TB (*22*). We next determined PI 16:0_19:0 levels in lung homogenates of these mice. During the whole course of infection PI 16:0_19:0 concentrations stayed at ~1.1×10^-2^ pmol/μg in IL-13^WT^ mice (Fig. 3D), a value which was below the level of uninfected mice of 1.75×10^-2^ pmol/μg (n=11, dotted line in Fig 3D), which may indicate a metabolic background obtained from mice housed in an animal facility. It should be noted that for *in vivo* samples the mass spectrometric detected 19:0 FA comprises besides TSA also other isomers representing a metabolic background from microbiota and nutritional sources (fig. S4). Levels of PI 16:0_19:0 in lung homogenates of IL-13^TG^ were comparable to those in IL-13^WT^ mice during the early time points of the infection (d42, d62 p.i.), however at d139 in parallel with a significant rise in bacterial burden the amount of PI 16:0_19:0 increased and correlated with the *Mtb* loads in tissue samples (Fig. 3D, data file S2). Taken together, the detection limit at which PI 16:0_19:0 in tissue samples can be specifically recognized beyond the metabolic background of 1.75×10^-2^ pmol/μg was estimated to 1.0×10^5^ bacteria for an extracted lung tissue homogenate (Fig. 3D). Of note, whereas classical CFU determination requires 3-4 weeks before growth of *Mtb* could be determined, targeted quantification of PI 16:0_19:0 enables us to quantify increased *Mtb* loads in susceptible mice within 24 hours. However, the threshold of PI 16:0_19:0 concentration as correlate of CFU is only applicable for mouse models susceptible to *Mtb* infection, where bacterial loads are relatively high.

### Quantitation of PI 16:0 19:0 and putative transesterification products in PBMCs from TB patients and healthy controls

Next, we investigated whether the detection of PI 16:0_19:0 could be of potential use in a clinical setting. Attempts to monitor PI 16:0_19:0 levels in plasma samples from healthy controls and TB patients were not successful, very likely due to differences in sampling time caused by circadian rhythm and nutritional status (*23, 24*). We then hypothesized that mycobacterial PI 16:0_19:0 may be detectable in PBMCs isolated from active tuberculosis patients as these cells may have taken up this molecule as they are constantly migrating in and out of granulomatous lesions during the immune response against the pathogen (*25, 26*).

First, we analyzed PMBCs of 39 healthy controls and determined a metabolic background in these samples of about 0.13 pmol/1×10^6^ cells (median, n=38, data file S3), where for one individual PI 16:0_19:0 was below detection limit. The same analysis of PBMC from TB patients at therapy start (t0) showed significantly increased PI 16:0_19:0 levels of 0.3 pmol/1×10^6^ cells (median, n=23) when compared to healthy controls (Fig. 4A, data file S4). Receiver operating characteristic (ROC) were determined in comparison to the healthy controls with an AUC = 0.784 and sensitivity of 69.7% and specificity of 79.0% at the optimal cut off value (table S1, fig. S9). From the cohort of TB patients, we then focused on the subset of patients that either completed anti-tuberculosis therapy (WHO outcome criteria, Fig. 4B, n=13) or were cured according to the TBnet outcome criteria (*2*), which involves a follow up of one year after therapy end (Fig. 4C, n=9). We compared the levels of PI 16:0_19:0 in PBMCs at t0 and at the end of therapy (tE, Fig. 4C, data file S5). Considering the WHO criteria, PI 16:0_19:0 levels decreased in 9/13 patients, stayed similar in 3/13 and increased in 1/13 patients when t0 levels were compared towards tE. With regard to TBnet criteria PI 16:0_19:0 levels decreased for 7/9 and remained similar for 2/9 patients (Fig. 4B, C). Taken together, with both outcome definitions, we observed that PI 16:0_19:0 levels in PBMCs of these patients were significantly reduced at the end of therapy when compared to therapy naive patients.

**Fig. 4.**
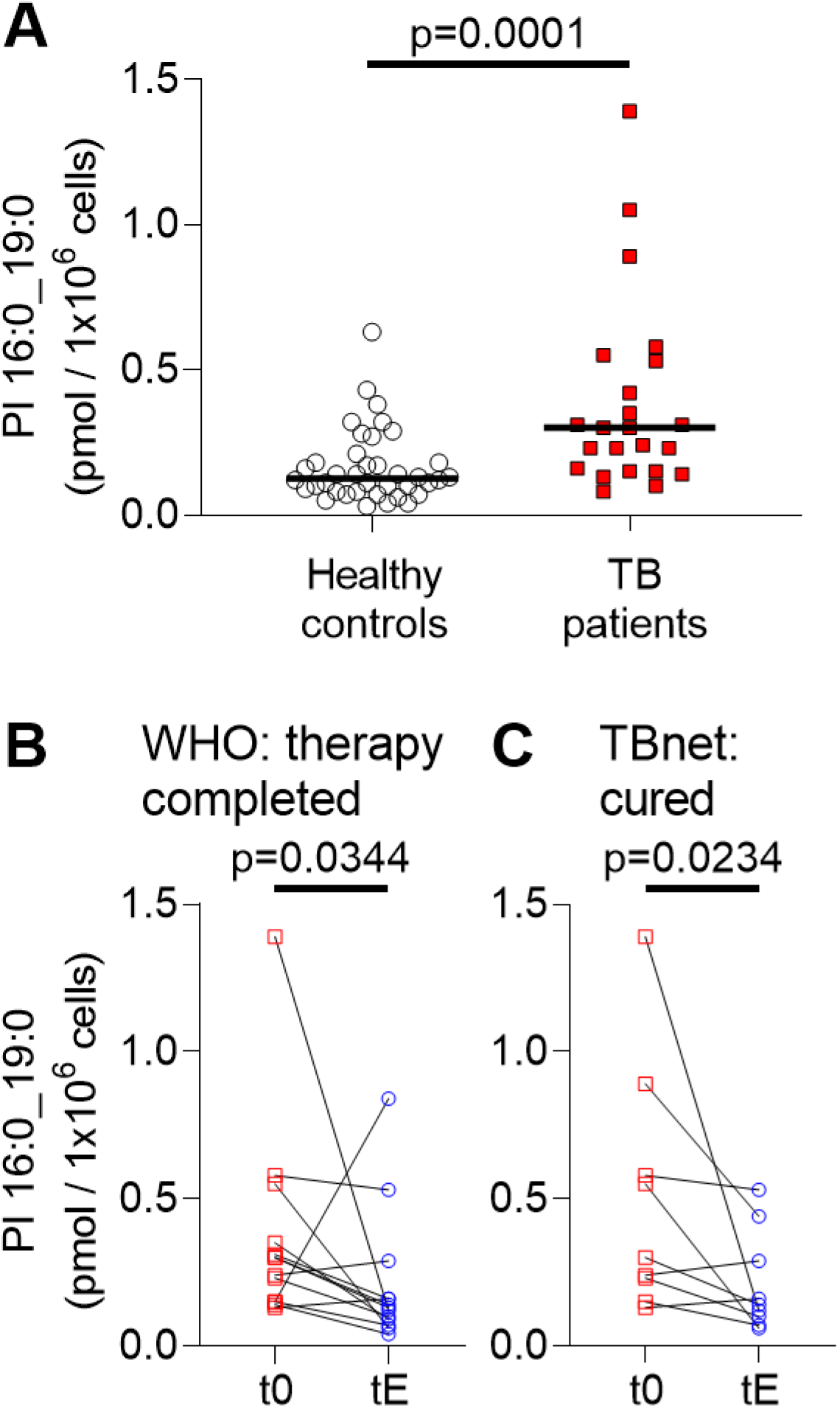
Amount of PI 16:0_19:0 in PBMCs of TB patients are indicative for disease status. **A)** Concentration of PI 16:0_19:0 in PBMCs of 39 healthy individuals (n=1 negative for PI 16:0_19:0) and 23 TB patients at therapy start (t0). Median is indicated with a black bar. Statistical significance was examined with Mann-Whitney test (individual data from patients and healthy controls is listed in data files S3, S4). **B)** PI 16:0_19:0 concentration at therapy start (t0) and at therapy end (tE) for 13 patients with the WHO defined outcome “therapy completed”. **C)** Comparison of PI 16:0_19:0 concentration for 9 patients at therapy start (t0) and therapy end (tE) that further fulfilled the outcome criteria “cured” following the TBnet criteria. Statistical examination for **B, C** was performed with a Wilcoxon matched-pairs signed rank test. All statistical tests were performed with GraphPad Prism 8.4.3. Data for comparison of TB patients at t0 and tE according to the outcome criteria definition are listed in data file S5.

We further hypothesized that host cells may have metabolized a certain amount of mycobacterial PI 16:0_19:0 via transesterification to form host cell lipids that contain TSA. Specifically, PI 19:0_20:4 caught our interest since it was detected with up to 20-fold higher amounts in PBMCs when compared to PI 16:0_19:0 and showed even slightly improved accuracy (fig. S10, table S1). This molecule is structurally very similar to the major abundant endogenous species PI 18:0_20:4 and might be produced with high transesterification rates. We further included seven additional PI species comprising FA 19:0 that were regularly detected in PBMCs into a marker panel (sum FA 19:0, table S2). For PBMC samples, the marker panel comprises a mix of PI isomers (fig. S4) where we expect that for TB patients increased contribution from TSA-containing PIs is quantifiable. A comparison of the overall amount of this maker panel between healthy controls and therapy naive TB patients showed such significant increase from a median of 5.1 pmol/1×10^6^ to 13.2 pmol/1×10^6^ cell, respectively (Fig. 5A, data files S3, S4). Furthermore, the accuracy for the comparison of TB patients and healthy controls were slightly better than for PI 16:0_19:0 alone with an AUC = 0.802 and sensitivity of 73.9%, and specificity of 82.1% (table S1, fig. S11). We then again focused on patients that were grouped either according to the WHO or the TBnet outcome definitions. The concentration levels of all FA 19:0 containing PIs in PBMCs showed a significant reduction at tE when compared with t0 (Fig. 5B, C, data file S5). In 9 out of 10 patients that were cured according to TBnet definition, the concentration of sum FA 19:0 was lower in PBMCs at tE when compared to t0 (Fig. 5C).

**Fig. 5.**
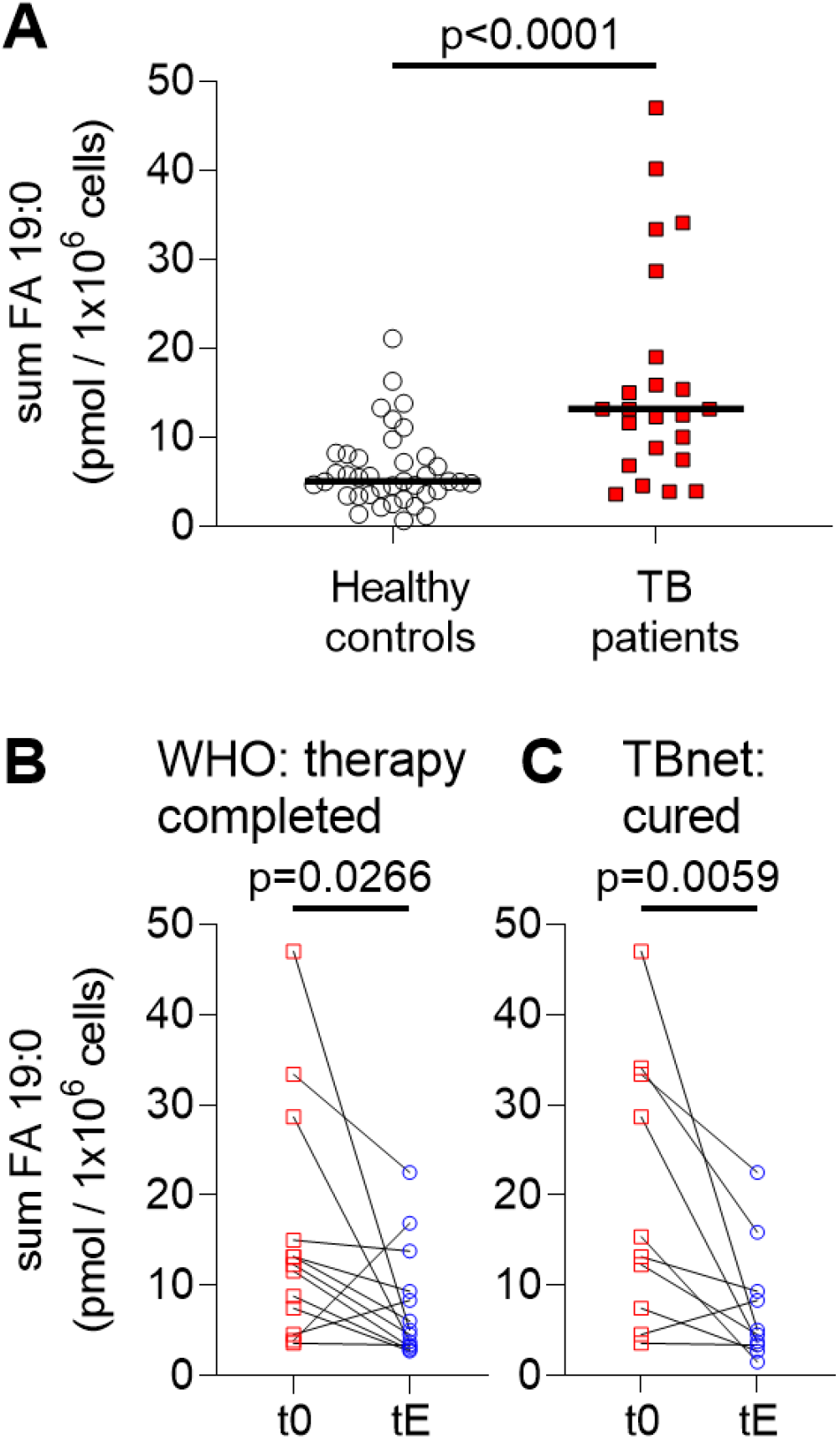
FA 19:0 containing PIs in PBMC correlate with therapy outcome for TB patients. **A)** Comparison of summed amounts of 9 FA 19:0 containing PIs in PBMCs (sum FA 19:0; table S2) of 39 healthy individuals with 23 TB patients at therapy start (t0). Median values are indicated with a black bar. Statistical significance was examined with Mann-Whitney test (data files S3, S4). **B)** Amounts of sum FA 19:0 at the therapy start (t0) and at therapy end (tE) for 13 patients with the WHO defined outcome “therapy completed”. **C)** Comparison of sum FA 19:0 for 10 patients at therapy start (t0) and therapy end (tE) that further fulfilled the outcome criteria “cured” following the TBnet criteria. Statistical examination for **B, C** was performed with a Wilcoxon matched-pairs signed rank test. All statistical tests were performed with GraphPad Prism 8.4.3. All data for paired statistical analyses according to TB outcome criteria are listed in data file S5.

## Discussion

We detected PI 16:0_19:0 (TSA) in 18 clinical MTBC isolates as abundant mycobacterial determinant representing different phylogenetic lineages. PI 16:0_19:0 quantifications correlated with CFU in broth cultures, cultures from murine macrophages and human neutrophiles, as well as in tissue samples of experimentally *Mtb* infected mice with a susceptible genetic background. Measuring PI 16:0_19:0 concentrations allowed to rapidly determine the mycobacterial burden within one day while CFU based enumeration requires up to 2 months. We further showed that biosynthesis of TSA by the pathogen and its incorporation into PIs but also other phosphoglycerolipids can become an interesting tool for metabolic studies (Fig. 6A). Furthermore, first proof of principle experiments using PBMCs from active TB patients showed enhanced levels of PI 16:0_19:0 and FA 19:0-containing PIs when compared to healthy controls and cured TB patients. We envisage that our study on the metabolic interaction of *Mtb* and its host, with focus on major abundant PIs containing FA 19:0, might encourage follow-up studies in the TB community (Fig. 6B). In the following, we discuss the open questions for the application of this lipid panel in translational research and clinical settings.

**Fig. 6.**
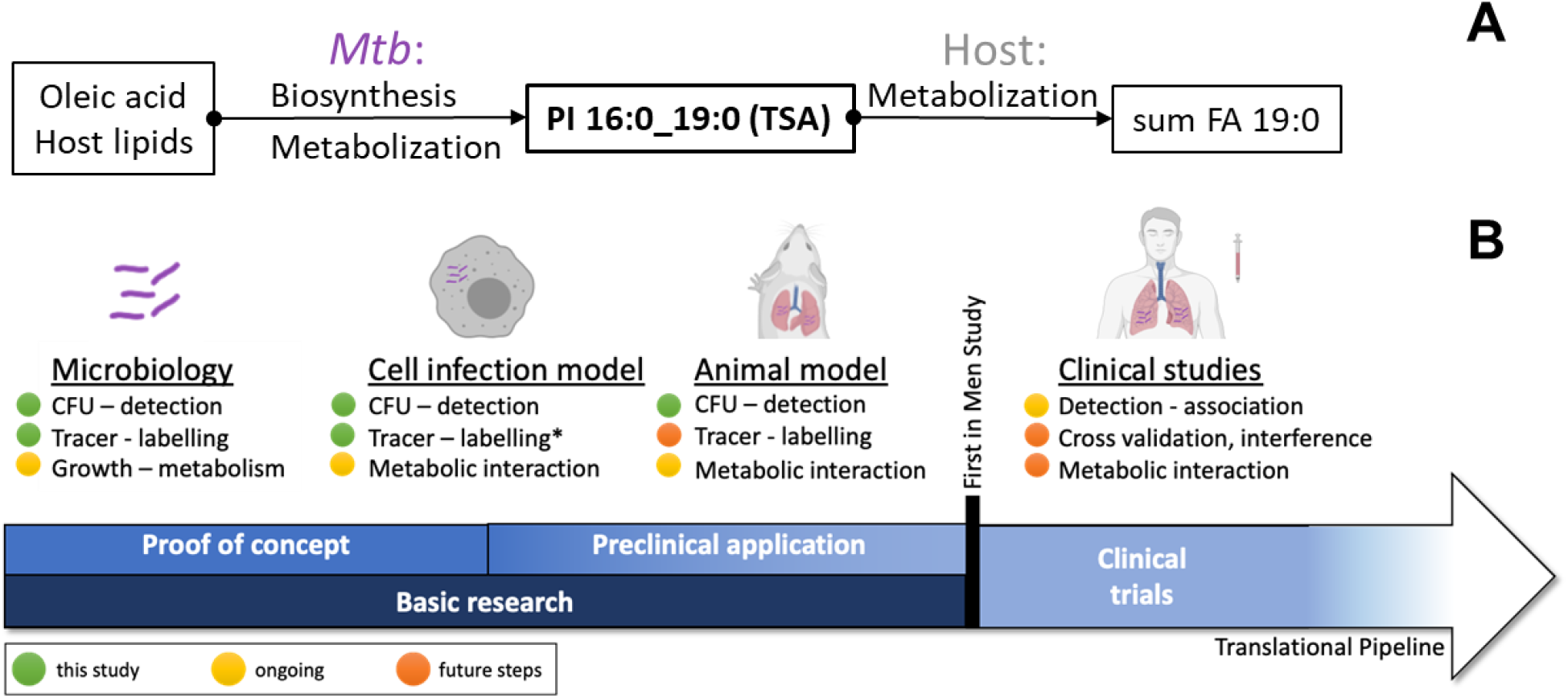
Implementation status for FA 19:0 (TSA)-containing mycobacterial PIs in translational TB research. **A)** TSA biosynthesis was observed from FA 18:1 (OA) as educt and host's complex lipids that contain OA by tracer analysis. The mycobacterial lipid PI 16:0_19:0 (TSA) itself was quantified in *in vitro* infection model system. Putative metabolization routes of TSA-containing PIs led to definition of the sum FA 19:0 marker panel. **B)** The FA 19:0 (TSA)-containing PIs of mycobacterial origin were tested in biological model systems along translational research pipeline. Studies on metabolization of PI 16:0_19:0 (TSA) in animal models are required to verify its application in the clinic. Further studies on interferences by other lung diseases and infectious diseases are required to validate TSA-containing PIs as marker for TB (* demonstrated in (*16*)).

### PI 16:0_19:0 (TSA) as marker for Mtb CFU and metabolic activity

In this study, we demonstrate that PI 16:0_19:0 can be used as a molecular marker of *Mtb* quantification in the background of complex sample matrices of mammalian origin. The direct link of PI 16:0_19:0 to the composition of the PM provides a solid basis for its usage for quantification of mycobacterial loads across the MTBC (Fig. 6A). Quantification of PIs is well established in lipidomics and appropriate internal standards for absolute quantitation are available (*27*). In addition, the analytical sensitivity for *Mtb* detection can be improved by targeted analysis using modern triple quadrupole instruments in combination with LC. This could improve detection limits that were in the current study approximately 100 CFU in bacterial culture systems and around 1,000 bacteria in cell culture systems. Tracer analysis using isotopically labelled OA will enable studying general aspects of the *Mtb* lipid metabolism. The incorporation of one ^12^C methyl group by *Mtb* to form ^12^C1^13^C18-19:0 (TSA) provides a very specific mass spectrometric tag to perform pulse-chase type of experiments in basic TB research. The enzymatic transformation of FA 18:1 (OA) to FA 19:0 (TSA) has been controversial discussed and two mechanisms have been proposed. Initially thought to be solely associated with Rv0447c, a two-step mechanism was suggested for *M. chlorophenolicum*, putatively catalyzed by proteins WP_048472120 and WP_048472121, orthologues of WP_003917236 and WP_003420415 (corresponding to Rv3719 and Rv3720 of H37Rv) (*28, 29*). We are convinced that further detailed analyses of the enzymatic transformation of FA 18:1 (OA) to FA 19:0 (TSA) in appropriate model systems will support studies on *Mtb* metabolism with regard to replication, dormancy, persistence and also reactivation.

Our understanding of the metabolization of PI 16:0_19:0 and TSA during interaction with the host into the metabolites described herein is incomplete. Most prominently, PI 19:0_20:4 as a surrogate mycobacterial marker suggests putative metabolization routes in the host for PI 16:0_19:0 (TSA) and/or other TSA-containing lipids via transesterification. However, basal levels for PI 19:0_20:4 and other FA 19:0 containing PIs were detected in non-infected murine lung tissues (data file S2), human neutrophils and PBMCs of healthy controls. This result indicates that interferences through nutritional and/or microbial factors need to be considered in future analyses. Finally, the metabolic steps responsible for the transformation of mycobacterial PI 16:0_19:0 or FA 19:0 (TSA)-containing lipids to PI 19:0_20:4, most likely linked to salvage pathways from lysosomes and phagosomal processing, are yet to be discovered (Fig. 6).

### Quantifying mycobacterial derived metabolites

Mycobacteria derived (glyco-)lipid markers like MAs (*12, 13*), LAM (*30, 31*), and also TSA (*32–34*) were already introduced for the diagnosis of active TB. Specifically, LAM assays were developed to identify TB cases in resource-limited settings. MAs were applied as discriminator in the clinical diagnosis of TB and might be used in drug efficacy analyses in mouse models (*12*). However, we assessed it critical that a fixed stoichiometry per bacteria cannot be expected for MAs as building block of the highly lipophilic outer membrane layer of the mycobacterial cell wall. MAs are mainly conjugated to the PGN-AG complex, but also covalently linked to trehalose to form TDM and trehalose monomycolate (TMM) and further present as free MAs (*6, 35, 36*). Therefore, MAs represent a dynamic class of cell wall molecules that cannot be assumed to be conserved in quantity, structure and composition during host pathogen interactions since they originate from several characteristic cell wall components that contain hundreds of molecular species. Specifically, TDM is expected to be generated in different quantities per bacteria as it is a virulence factor involved in hindering phago-lysosome fusion (*37–39*). Indeed, it was demonstrated that the amounts of TDM are enhanced in mycobacteria during the intracellular life phase in macrophages (*40*). While the complex regulation of MAs by *Mtb* is of utmost interest for the basic understanding of host pathogen interaction, we found that the degree of structural complexity, especially caused by various modifications like cyclopropane rings, methyl branches, ketones and methoxy groups (*35, 41*) and unknown metabolic pathways in the host are discouraging for further marker development. Even more important, it was shown that the MA profiles are strain dependent (*12*) and we aimed to utilize marker molecules that are present in the entire MTBC.

FA 19:0 (TSA) as marker has two major advantages: it is present in large amounts in all prototype isolates of the MTBC we investigated so far and its quantitation can directly be performed by GC-MS (*34, 42*). However, as chemical building block it is present in a number of important mycobacterial lipids like the major abundant PL classes PE and PI as well as PIMs (*43*), LM and LAM (*44*). Specifically, the latter complex glycoconjugates are reported as virulence factors. Glycolipids are either inserted in the plasma membrane stretching through the cell wall to the outer surface or are linked to the outer membrane (*45*). In analogy to MAs, it is likely that the amount of FA 19:0 (TSA) per bacteria varies depending on environmental condition, metabolic status and host pathogen interaction. In contrast, the PI composition of the PM was well conserved between all investigated MTBC strains. Thus, our focus on FA 19:0 (TSA)-containing PIs and its major abundant species present in the PM of *Mtb* could provide a robust correlation to the mycobacterial burden in experimental model systems (Fig. 6A).

### Perspectives for mycobacterial derived lipid biomarker in clinical application

Commercially available PCR based test systems such as the GeneXpert MTB/RIF ultra (Cepheid, Sunnyvale, Ca., USA) are very sensitive with regard to the detection of *Mtb,* but are most probably not suitable for therapy response assessment due to extended mycobacterial DNA stability even after the death of the bacteria (*46, 47*). However, PCR based methods are able to predict drugresistance for most anti-TB agents, which is not possible based on lipids (*48*). Currently, culture remains the gold standard for the diagnosis of TB and the assessment of bacterial burden despite being slow. In the present study, we performed first analyses with regard to the presence of PI 16:0_19:0 and/or other FA 19:0 -containing lipids in TB patient PBMCs. Future studies evaluating its usability as blood-based diagnostic tool may discover interference with non-tuberculous mycobacteria (NTM) and other *Actinomycetales*. In a future perspective we nevertheless could envisage that quantitation of PI 16:0_19:0 may allow a more rapid estimation for *Mtb* burden even during TB therapy, which could be potentially expanded to sputum samples as well. In order to get to this point, detailed longitudinal studies of patient samples taken at different time points of therapy are required to foster such an approach (Fig. 6B).

There is ample evidence for the strong dependency of the plasma lipidome composition on sampling time caused by circadian rhythm and nutritional status (*23, 24*). In addition, several factors related to the patient's health and to local healthcare settings may prevent a timely blood sampling at the same time point. That is why we focused our analytical developments on the quantitation of lipid markers in PBMCs. By this strategy, we aimed to minimize the variation in lipid composition caused by the metabolic state of patients. With regard to this notion, our findings are encouraging to further explore mycobacterial lipid biomarkers in clinical application based on well-established PBMC isolation protocols.

In the presented study, we did not address in detail whether an interaction between response profiles of FA 19:0 -containing PIs and PBMC cell populations exists. Further studies are required to evaluate such correlations and whether performance of the marker could be improved by focusing on specific subsets of immune cells such as professional phagocytes. We are confident that the presented approach for the clinical sample preparation could also be performed in low-resource countries when sample storage and transport would follow general sample handling guidelines. Specifically, an early addition of methanol and the antioxidant butylhydroxytoluol (BHT) to PBMCs should sufficiently block inherent enzymatic activities and oxidation. As diagnostic markers, PI molecules are chemically stable enough to establish recruitment and sampling strategies, which make the transfer of patient samples to dedicated MS facilities possible.

## Materials and Methods

### Mycobacteria

MTBC strains (Table 1) were grown in Middlebrook 7H9 broth (Difco, Detroit, USA) supplemented with 10% Middlebrook oleic acid albumin-dextrose-catalase (OADC) enrichment medium (Life technologies, Gaithersburg, USA), 0.2% glycerol and 0.05% Tween 80 in 490 cm^2^ Corning^®^roller bottles. Cultures were harvested at mid log phase (OD600 = 0.3 – 0.6), aliquoted and frozen at −80 °C.

### Cultivation and lipidomics analysis of MTBC clinical isolates

Frozen aliquots (OD_600_ = 0.3 – 0.6) of clinical isolates were thawed and homogenized by transferring bacterial suspension through a 1 mL-syringe. A total of 300 – 500 μL of bacterial culture were used for inoculation in a final volume of 10 mL. Cultures were incubated at 37 °C and 5% CO2 for 7 – 10 days. Contamination controls on blood agar and Ziehl – Neelsen staining were performed repeatedly. The bacterial load was determined by CFU analysis on 7H10 agar plates containing 10% bovine serum and 0.5% glycerol. The analytical strategy to profile mycobacterial lipids is summarized in Supplementary Materials and Methods (fig. S11). Briefly, bacterial pellets of 5.0×10^8^ bacteria were transferred into glass tubes containing 2 mL petroleum ether (bp 60-80 °C) and 4 mL methanol for inactivation of bacteria. Only the methanolic phase was analyzed with LC-MS using an 1100 HPLC system (Agilent, Waldbronn, Germany) coupled to an Orbitrap Q Exactive Plus (Thermo Fisher Scientific, Bremen, Germany). 100 μL of the methanolic phase were dried under vacuum and reconstituted in 100 μL of CHCl3/MeOH containing 0.1% ammonium acetate 86/13 (v/v) and 5 μL were used for injection. Separation of lipids was performed on a 150 mm BETASIL Diol-100 column with a particle size of 5 μm and 0.32 mm inner diameter (Thermo Fisher Scientific, Bremen, Germany). Detailed information for LC-MS data acquisition is given in Supplementary Materials and Methods.

### Detection of PI 16:0_19:0 as surrogate for CFU determination

Aliquots of mCherry expressing *Mtb* H37Rv (OD_600_ = 0.4) were thawed and centrifuged (4,200 x *g*, 10 min, 4 °C). After resuspending the pellet in PBS, defined amounts ranging from 1.0×10^2^ to 1.0×10^7^ bacteria were transferred into tubes and again centrifuged (4,200 x *g*, 10 min, 4 °C). Pelleted bacteria were then agitated in 400 μL methanol for 16 h prior to lipid extraction and shotgun lipidomics analysis.

### Detection of Mtb in artificial sputum with targeted analysis of PI 16:0_19:0

Artificial sputum was prepared according to a customized protocol (*49*) and spiked with known CFU of *Mtb* H37Rv and background flora composed of fungal and bacterial pellets. Pellets were mixed with 1 mL MeOH to kill bacteria in the biosafety laboratory level 3 and transferred to the chemical laboratory. The samples were dried down in a SpeedVac vacuum concentrator (Thermo Fisher Scientific, Waltham, US) before they were resuspended in 50 μL water. The suspension was thoroughly mixed and shock-frozen in liquid nitrogen. Afterwards, samples were transferred into an ultrasonic bath (Sonorex, Bandelin, Berlin, Germany) filled with ice-cooled water. Samples were sonicated for 1-2 min until thawing was complete. This freeze/thaw cycle was repeated three times. Then the modified MTBE protocol described below was performed. The organic phase was dried down and dissolved directly in 100 μL MS ESI solution consisting of CHCl3/MeOH/isopropanol 1/2/4 (v/v/v) with 0.05 mM ammonium chloride for shotgun lipidome analysis.

### Isotopic labelling of Mtb phospholipids

2.0×10^6^ GFP expressing *Mtb* bacteria H37Rv were cultured in a microtiter based format with a volume of 100 μL (*50*) in 7H9 medium containing 0.5% BSA, 0.085% NaCl, 0.0003% Catalase and 0.04% Na-Acetate. Culture media were supplemented with natural isotope distribution (^12^C) or completely ^13^C-labeled oleic acid (^12^C: O1383, Sigma Aldrich, Taufkirchen, Germany; ^13^C: CLM-460-0 (98%), Cambridge Isotope Laboratories, Tewksbury, US). After 7 days of culture, bacterial suspensions were centrifuged at 4,200 x *g* for 10 min at 4 °C, followed by removal of the supernatants. The pellet was resuspended and agitated in 500 μL methanol for 2 h at RT prior to lipid extraction and shotgun lipdomics.

### Generation of bone-marrow derived macrophages (BMDM) and infection with Mtb

Murine bone marrow derived macrophages (BMDM) were isolated as described earlier (details are given in Supplementary Materials and Methods) (*51, 52*). For infection of BMDM, frozen aliquots of *Mtb* H37Rv were thawed, centrifuged (2,300 x *g*, 10 min) and bacteria in cell culture medium carefully homogenized by use of a 26-gauge syringe needle. Subsequently, cells were infected with a MOI of 1 and were incubated for 4 h (37 °C, 5% CO2), followed by extensive washing with Hanks Buffered Salt Solution (HBSS, Sigma Aldrich, Taufkirchen, Germany) in order to remove extracellular bacteria. Cells were cultured for a total of 72 h as previously described (*53*). Afterwards, supernatants were discarded, and cells washed once with PBS (37 °C). Subsequently, 100 μL of ice-cold water were added, and cells were incubated on ice for 30 min. Cells were detached by use of a cell scraper and transferred into suitable tubes (Eppendorf, Hamburg, Germany) containing 270 μL of MeOH (hypergrade for liquid chromatography, Merck, Darmstadt, Germany).

### Isolation and infection of peripheral human neutrophils

Human neutrophils were isolated from peripheral blood of healthy volunteers (two female and three male of normal weight) aged 30 – 40 years. For the isolation of peripheral neutrophils, granulocytes were purified by density centrifugation for 25 min at RT at 800 x *g* through a Histopaque 1199 gradient (Sigma Aldrich, Taufkirchen, Germany). A discontinuous Percoll^®^ gradient (Sigma Aldrich, Taufkirchen, Germany) was subsequently used to isolate polymorphonuclear leukocytes (PMN). The cells were seeded at 5.0×10^6^ cells/mL in RPMI medium (PAA, Traun, Austria) supplemented with 2 mM L-glutamin and 10% heat inactivated, autologous donor serum. For infection, an*Mtb* H37Rv suspension was pelleted and washed twice in RPMI. Upon resuspension in RPMI, mycobacteria were opsonized with fresh autologous donor serum for 30 min at RT. PMN were infected with an MOI of 3 and incubated for 6 h at 37 °C with 5% CO2. To assess the infection rate of the PMN culture, CFU were determined as described in Supplementary Materials and Methods. For targeted lipid analysis, the second aliquot of 1 mL cell culture was washed twice with PBS and finally harvested using a centrifugation for 10 min at 15,000 x *g* at 4 °C. Cell pellets were resuspended in 50 μL PBS and transferred into tubes containing 270 μL methanol + 0.184% BHT (w/v) and incubated over night at 4 °C to inactivate the bacteria. Lipid extraction was afterwards continued according to the extraction protocol described below.

### Mycobacterial infection of mice

Interleukin (IL)-13-overexpressing (IL-13^TG^) and wild-type littermates (IL-13^WT^) on a BALB/c genetic background (*20*) were bred under specific-pathogen free conditions. For infection experiments, 8- to 12-week old mice were transferred to the BSL3 facility at the Research Center Borstel and kept in individually ventilated cages. Mice were infected with a low dose of appr. 100 CFU of *Mtb* H37Rv via aerosol and bacterial loads in lungs were determined at 42, 62 and 139 days after infection as described earlier (*54*). To enable standardized lipid extraction of lung tissues, the earlier described method (*27*) was modified (for details see Supplementary Materials and Methods). Briefly, tissue homogenate aliquots containing 0.012 g wet weight were extracted and the total protein content was determined after MTBE extraction. Normalized CFU for each sample were calculated in concordance to the volume of each tissues homogenate aliquot used for lipid extraction (data file S2). Lipid extraction was performed according to a customized MTBE-based extraction as described earlier ((*27*), see details in Supplementary Material and Methods).

### Description of patient cohort

Patients with pulmonary TB (non-M/XDR-TB and M/XDR-TB) were enrolled at five respiratory care centers in Germany as previously described (*48, 55*). The study inclusion criterion was active pulmonary TB confirmed by a molecular test (GeneXpert^®^, Cepheid, Sunnyvale, USA). Written informed consent was obtained from all study patients. Blood for PBMC analysis was sampled before therapy start (t0: defined as time point 0-14 days post diagnosis), when this was clinically not possible the number of days after the ideal baseline sampling was recorded (data file S4, S5). Subsequently, samples were taken at predefined time points throughout drug-treatment. Patient therapy outcomes were assessed using the TBnet simplified outcome criteria (*2*) and WHO defined criteria (*56*).

### Healthy cohort of the Research Center Borstel

39 healthy adult Caucasian volunteers were enrolled at the Center for Clinical Studies, Research Center Borstel. Due to defined exclusion criteria no subject was treated with systemic steroids during the last month, antibiotics in the previous two months, suffered from Asthma, COPD (stage GOLD III/IV), airway infections during the previous month, any immunosuppression, diabetes mellitus, previous or active tuberculosis, was pregnant or breast feeding.

### Ethic statements

Animal experiments were approved by the Ministry of Energy, Agriculture, Environment, Nature and Digitization of the state of Schleswig-Holstein, Germany (Approval no. 69-6/16).

Study approval for human samples from active TB patients (AZ 12-233) and healthy volunteers (AZ 15-194) was granted by the Ethics Committee of the University of Lübeck.

For isolation of human neutrophils, written consent approving and authorizing the use of the volunteers’ material for research purposes was obtained from all donors. The Ethics Committee of the University of Lübeck approved isolation of and experimentation with human peripheral blood cells (12-202A and 19-071).

### Isolation of peripheral blood mononuclear cells samples

Blood for isolation of mononuclear cells (PBMC) was sampled in CPT™ vacutainer (BD, Heidelberg, Germany). PBMCs were isolated according to manufacturer’s instructions from heparinized whole blood. PBMC were harvested from the interface, washed twice in PBS (Biochrom, Berlin, Germany) (250 x *g*, 15 min, 4 °C) and counted (Casy2, Schärfe System, Reutlingen, Germany). Subsequently cell pellets were stored at −80 °C prior to MS analysis.

### Targeted quantitation of TSA-containing phosphatidylinositols of PBMC

PBMC pellets containing either 1×10^6^, 2×10^6^ or 5×10^6^ cells were washed with PBS and afterwards taken up in 50 μL water. After thoroughly mixing of the samples an internal standard mix was added (Supplementary Materials and Methods, table S3). All samples were extracted following a customized methyl-*tert*-butyl ether (MTBE) protocol (*27, 57*). Briefly, first 270 μL MeOH containing 3% acetic acid were added. Second, 1 mL MTBE was added to the mixture resulting in a one-phase system that was incubated at RT for one hour under constant shaking. For separation of lipids from hydrophilic components the phase separation was induced by addition of 250 μL water. After an additional incubation period of 5 min, the mixture was centrifuged for 2 min at RT at 13,400 x *g*. 800 μL of the upper organic phase were transferred into a new tube. The samples were extracted a second time under the same conditions. The combined organic phase was dried in a vacuum concentrator and subsequently dissolved in 50 μL storage solution (CHCl3:MeOH:H2O, 60/30/4.5, v/v/v). Lipid extracts were stored until analysis at −80 °C. Shotgun lipidomics was performed as reported earlier (*27*) using the TriVersa NanoMate (Advion, Ithaca, USA) as nanoelectrospray coupled to an Orbitrap Q Exactive Plus (Thermo Fisher Scientific, Bremen, Germany) mass spectrometer. A targeted analysis was performed over a time period of 5 min in the negative ion mode for PIs listed in Supplementary Materials and Methods table S2. Lipid identification was performed with LipidXplorer (*58, 59*). Lipid annotation was performed according to the fatty acyl / alkyl level as described earlier ((*60*), see details in Supplementary Materials and Methods, fig. S4). Quantities of TSA-containing PIs were calculated in relation to the response of the FA 18:1 (d7) fragment ion of the internal standard PI 15:0-18:1(d7) (828.5625 -> 288.2925).

## Acknowledgments

The authors acknowledge the excellent technical assistance of Birgit Kullmann, Romina Pritzkow, Franziska Daduna, Jessica Hofmeister and RCB blood donor service. We thank Andrea Glaewe, Lenka Krabbe, and Johanna Doehling for recruiting healthy volunteers and collecting samples. We acknowledge the excellent technical assistance of Doreen Beyer, Carolin Golin, Svenja Goldenbaum, and Alexandra Hölscher. We are also grateful to Anja Walter and Marion Schuldt for supplying and cleaning the lab and to Ilka Monath, Christine Keller, Sarah Vieten and Gerhard Schultheiß for organizing the animal facilities in Borstel and Kiel. We thank Prof. Dr. S. Niemann for critical review and discussion on the manuscript. The BioMaterialBank Nord is supported by the German Center for Lung Research and member of popgen 2.0 network (P2N). Fig. 6 was created using icons from BioRender.com.

## Author contributions

JH, NR and DS conceived and designed the study. NZ, HK, FW, LL, MW and VS performed lipidomics experiments and contributed to the data analysis. JB and LL performed cell culture experiments. FW and CHö performed mice experiments. NR, SH, SM performed mycobacterial cultivation and selected strains of MTBC. KK designed the bacteriological test system for sputum analysis and HK performed the lipid analysis. Recruitment and sample collection for TB patients was organized by JH, BK and CH. Recruitment and sample collection for healthy individual was managed by CHe and KD. Collection and evaluation of medical data was overseen by PSC, BK, CHe, KG, MR and JH. JB, JH, FW, NG, NR and DS drafted the manuscript. All authors edited and commented on the final manuscript.

## Data and materials availability

All data associated with this study are present in the paper or the Supplementary Materials.

## Supplementary Materials

### Methods

#### Extraction and LC-MS of amphiphilic mycobacterial lipids

Mycobacteria were grown in 7H9 media (BD Middlebrook 7H9 Broth) containing 10% OADC, 0.2% glycerol and 0.05% Tween 80 as described in the main text. Samples were washed twice with PBS and aliquots corresponding to 5.0×10^8^ bacteria in 1 mL PBS were transferred into glass tubes containing 2 mL petroleum ether (bp 60-80 °C) and 4 mL methanol (LC-MS grade) for inactivation of bacteria. After 15 min shaking at RT in a rotator, samples were centrifuged for 10 min at 400 x *g*. The upper petroleum ether phase was transferred to another vial and the methanolic phase was re-extracted one more time with 2 mL petroleum ether (fig. S12). The methanolic phase was separated from the intermediate layer and stored at −20 °C until analysis.

#### LC-MS analysis of the methanolic phase of the mycobacterial lipid extraction

Prior to LC-MS analysis, 100 μL of the methanolic phase were dried in vacuum and reconstituted in 100 μL of CHCl3/Methanol + 0.1% ammonium acetate 86/13 (v/v) and sealed using aluminum foil. An Agilent 1100 HPLC system (Agilent, Waldbronn, Germany) was used for the separation of the phospholipids on a 150 mm BETASIL Diol-100 column with a particle size of 5 μm (Thermo Fisher, Bremen, Germany) and 0.32 mm inner diameter using 5 μL injection volumes. The solvents, gradient profile and flow rates are summarized in table S4.

The spectrometric analyses were performed on a Q Exactive Plus instrument (Thermo Fisher Scientific, Bremen). For phospholipid analysis, full scan MS data (*m/z* 400-1800) were acquired in the first 18 minutes of the LC-run using positive and negative ion mode switching. Instrumental parameters were as follows: source temperature at 250 °C, −3.0 kV and 4.0 kV ionization voltage for negative and positive ion mode analysis, respectively. Resolution was set to 280 k, ACG target value was set at 3.0×10^6^ and RF value was at 100%. The sheath gas was set to 5 L/min, aux and sweep gas were set to 0 L/min. Data acquisition was performed using the Xcalibur 2.4 software (Thermo Fisher Scientific, Bremen).

#### Data Processing

LC-MS data files were sliced in raw files containing only positive or negative mass spectra using Thermo’s Slicer, Version 1.3. Raw files were converted to mzml using MSConvert (*67*) and files of the negative ion mode were analyzed using LipidXplorer. Lipids were assigned by their monoisotopic masses with a mass accuracy better than 5 ppm using LipidXplorer (*59*). Import settings and MFQL scripts will be made available at lifs.isas.de.

#### Generation of bone-marrow derived macrophages (BMDM)

Naval Medical Research Institute (NMRI) *Wnt6* gene-deficient and *Wnt6* wild-type mice (*68*) were raised and maintained under specific pathogen-free conditions. In order to generate BMDMs, mice were euthanized and femora and tibia were washed with ice-cold Dulbecco's Modified Eagle Medium (DMEM). Isolated bone-marrow cells were seeded on Nunclon Delta cell culture dishes (Thermo Fisher Scientific, Waltham, USA), and cultivated in macrophage cell culture medium (10 mM HEPES, 1 mM sodium pyruvate, 2 mM glutamine, 10% of heat-inactivated fetal calf serum (FCS; Biochrom, Berlin, Germany) in DMEM) containing 50 ng/mL macrophage colony stimulating factor (M-CSF; Bio-Techne, Minneapolis, USA). After 24 h, non-adhered cells were collected, seeded on cell culture dishes (Sarstedt, Nümbrecht, Germany) and cultivated for 7 days. Cells were incubated in macrophage cell culture medium overnight before proceeding further.

#### CFU determination in PMN culture

After incubation of *Mtb* infected PMN for 6 h at 37 °C with 5% CO2, one aliquot of 100 μL of the PMN culture was transferred to a new plate. Afterwards, the cell culture was centrifuged (500 x *g,* 5 min, RT) and the supernatant was discarded to remove non-phagocytized bacteria. The cells were lysed in 0.5% TritonX-100 in PBS and a 10-fold serial dilution in 0.05% Tween80 in PBS was prepared and plated on Middlebrook 7H11 agar plates. After incubation for 3-4 weeks at 37 °C colonies were counted and bacterial load per mL was calculated.

#### Lipid analysis of mice lung tissue homogenates and protein content determination

The lung wet weight of each tissue sample was noted before homogenization. For all extraction aliquots of each lung homogenate were inactivated in 80% methanol. For the normalization of the extraction procedure a volume corresponding 0.012 g of lung tissue was transferred in a new vial, dried in a SpeedVac and afterwards resuspended in 20 μL 50 mM ammonium acetate (data file S2). The lipid extraction was performed with a customized MTBE method as described earlier (*27*). Then 270 μL methanol, containing 3% acetic acid, were added. After vortexing the mixture, 30 μL of the internal standard solution (Avanti, SPLASH^®^ Lipidomics^®^ Mass Spec Standard, 330707) and 1 mL MTBE were added. The solution was incubated for one hour at RT with continuous shaking at 600 rpm (Eppendorf, MixMate). For phase separation 250 μL of water was added and subsequently incubated for 10 min at RT with continuous shaking at 1300 rpm. After incubation samples were centrifuged for 10 min at 15,000 x *g* and then the upper phase was collected in a separate tube. The lower phase was re-extracted with 400 μL theoretical upper phase (as described in (*57*)), vortexted and incubated for 20 min at RT with continuous shaking (1300 rpm). The solution was once more centrifuged as described above. The resulting upper phase was combined and the re-extraction step with the lower phase was repeated. In the end, the combined upper phase was dried in a SpeedVac. The dried extracts were dissolved in a mixture of chloroform, methanol and water (60/30/4.5; v/v/v) and stored at −80 °C. Shotgun lipidomics was performed on Q Exactive (Thermo Fisher Scientific, Bremen, Germany) as described earlier.

The water phases and non-extractable residue left after MTBE extraction were dried under vacuum. The sample pellet is resuspended in a 150 μL sample buffer (1 g SDS and 6.25 mL Tris/HCl (0.5 M, pH 6.8) filled to 100 mL with water). Samples are mixed well and are incubated for 15 min in a shaker (650 rpm) followed by a short centrifugation (2 min at 16,000 x g) step to have all sample located at the bottom of the vial. Protein quantification was performed with the Pierce^TM^ BCA^TM^ Protein-Assay Kit (Thermo Fisher Scientific, Waltham, US) according to the vendor's protocol.

#### ROC analysis

R version 3.4.0.2 was used to perform ROC analyses using the packages cutpointr and pROC to determine and visualize test performance of the parameters. Additionally, false positive and false negative rates were calculated based on the optimal cut-off classification.

#### Lipid annotation

The targeted MS^2^ enables to identify lipids until the level of fatty acid isomers. As characteristics ions for the PI class, we utilized *m/z* 241.01. However, as quantifier ion the acyl anion of the FA 19:0 (TSA) was utilized. For the annotation convention we followed the guidelines of Liebisch et al. (*60*). In this study we determined isomers on three levels:

1. bond type high resolution; PI 35:0
2. fatty acyl; PI 16:0_19:0
3. fatty acyl (*); PI 16:0_19:0 (TSA)

These levels are also depicted in view of the certainty of the structural assignment of possible isomers in fig. S4. For level 3, as shown in (Fig. 1, 2), we cannot exclude *sn*-position isomerism and determine isomeric purity. However, because of the utilization of ^13^C18-labeled OA and the methylation performed by *Mtb,* we are certain of the detection of TSA.

### Supplementary Figures

**Fig. S1.**
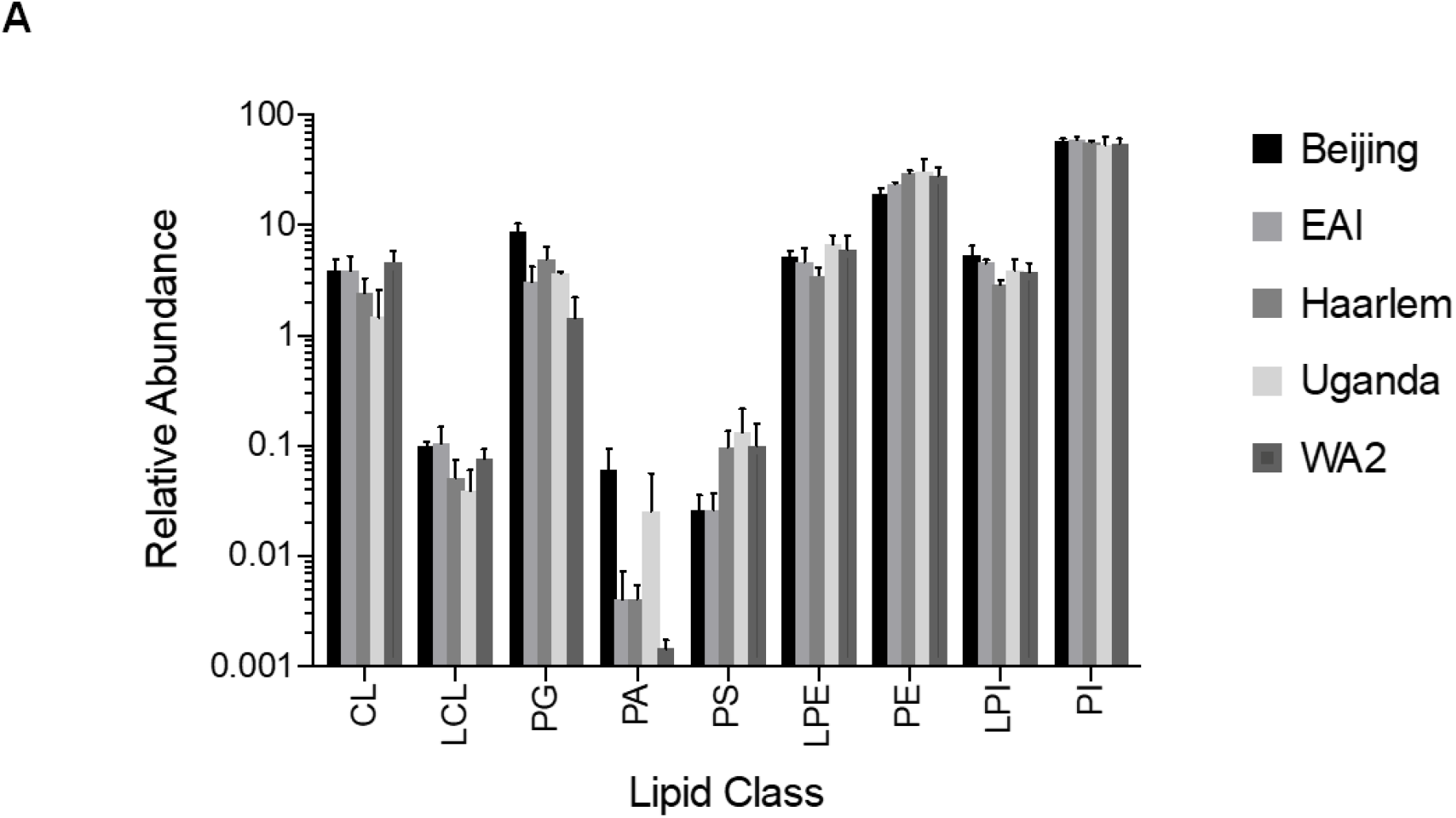
Overall composition of major abundant phospholipid classes of mycobacterial isolates. Lipid class distribution for all listed mycobacterial genotypes. Relative abundance represents the average of all strains of a genotype (**Table 1**). Relative abundances of lipids was determined by top down lipidomics approach using internal standards 4ME 16:0 Diether PE (1,2-di-O-phytanyl-*sn*-glycero-3-phosphoethanolamine) and PS 17:0/17:0 (both standards from Avanti Polar Lipids, Alabaster, US) for normalization of peak intensities. CL – cardiolipin; LCL – lysocardiolipin; PG – phosphatidylglycerol; PS, phosphatidic acid; PS – phosphatidylserine; LPE – lysophosphatidylethanolamine; PE – phosphatidylethanolamine; LPI – lysophosphatidylinositol; PI – phosphatidylinositol.

**Fig. S2.**
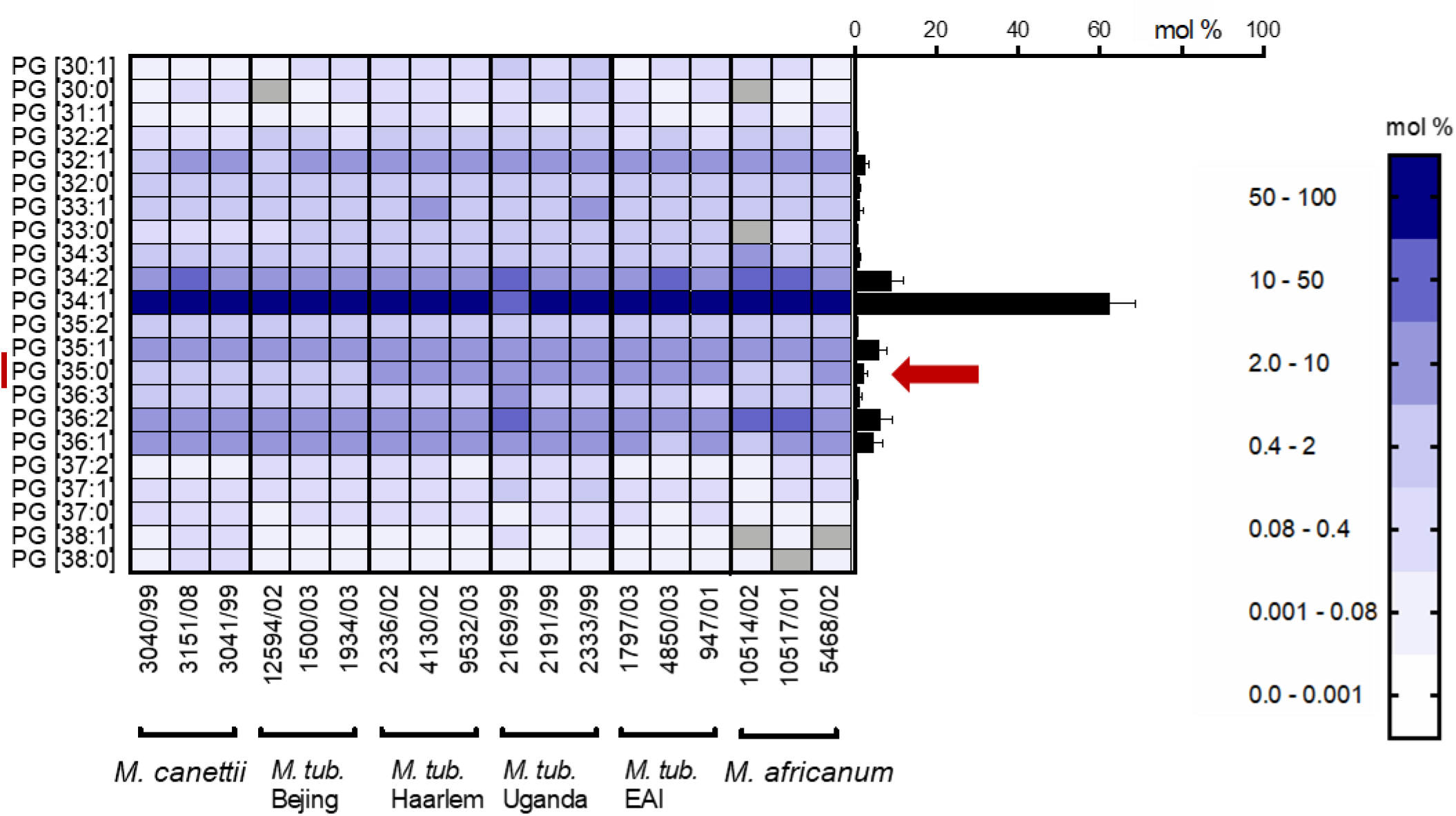
Lipid species distribution of major abundant mycobacterial phosphatidylglycerol (PG). Profiles of clinical isolates were determined from at least two independent isolations. Average profiles over all strains is represented in the right bar graph (Error bars represent one SD).

**Fig. S3.**
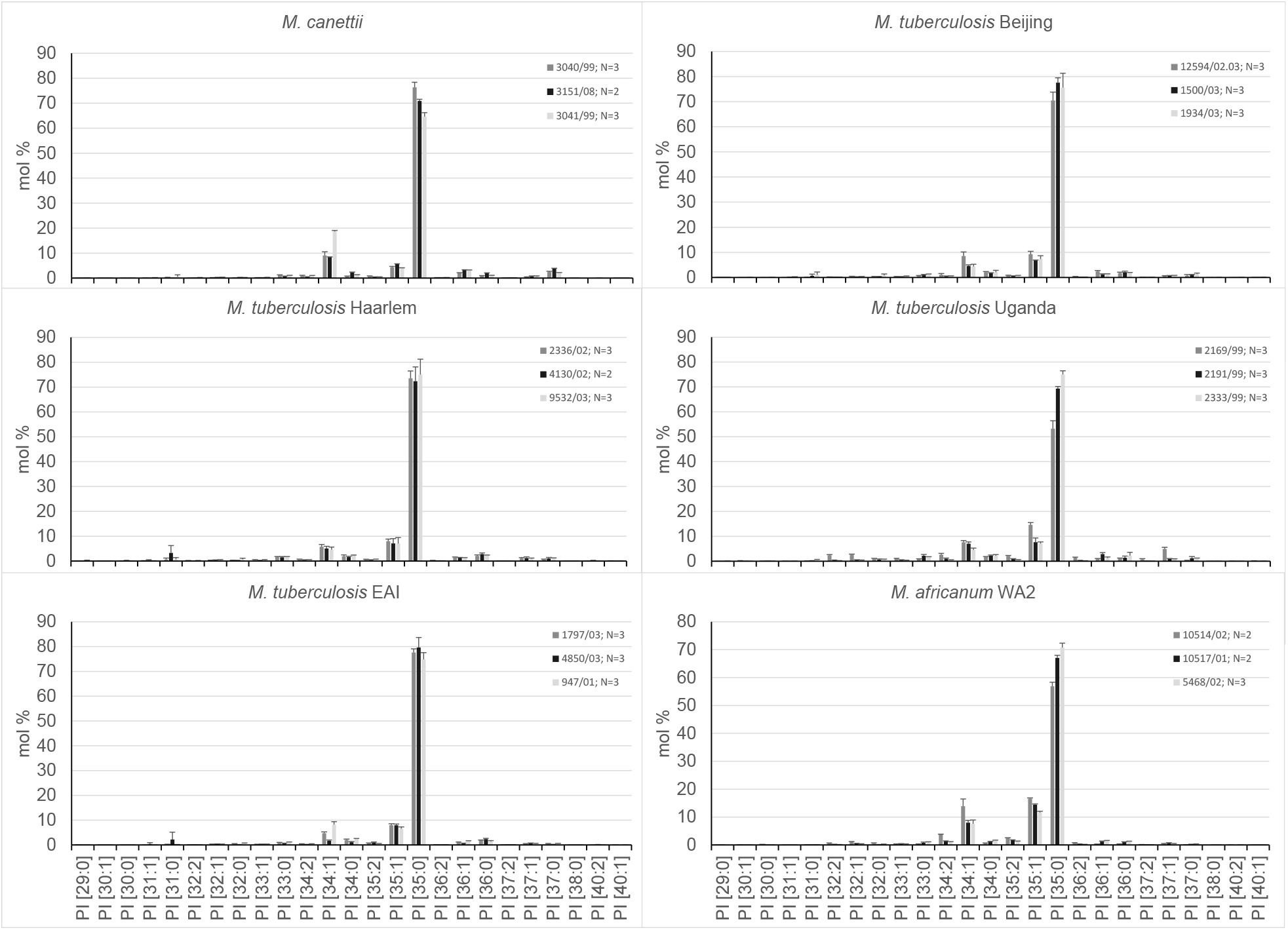
Distribution of PI species for 18 clinical MTBC isolates.

**Fig. S4.**
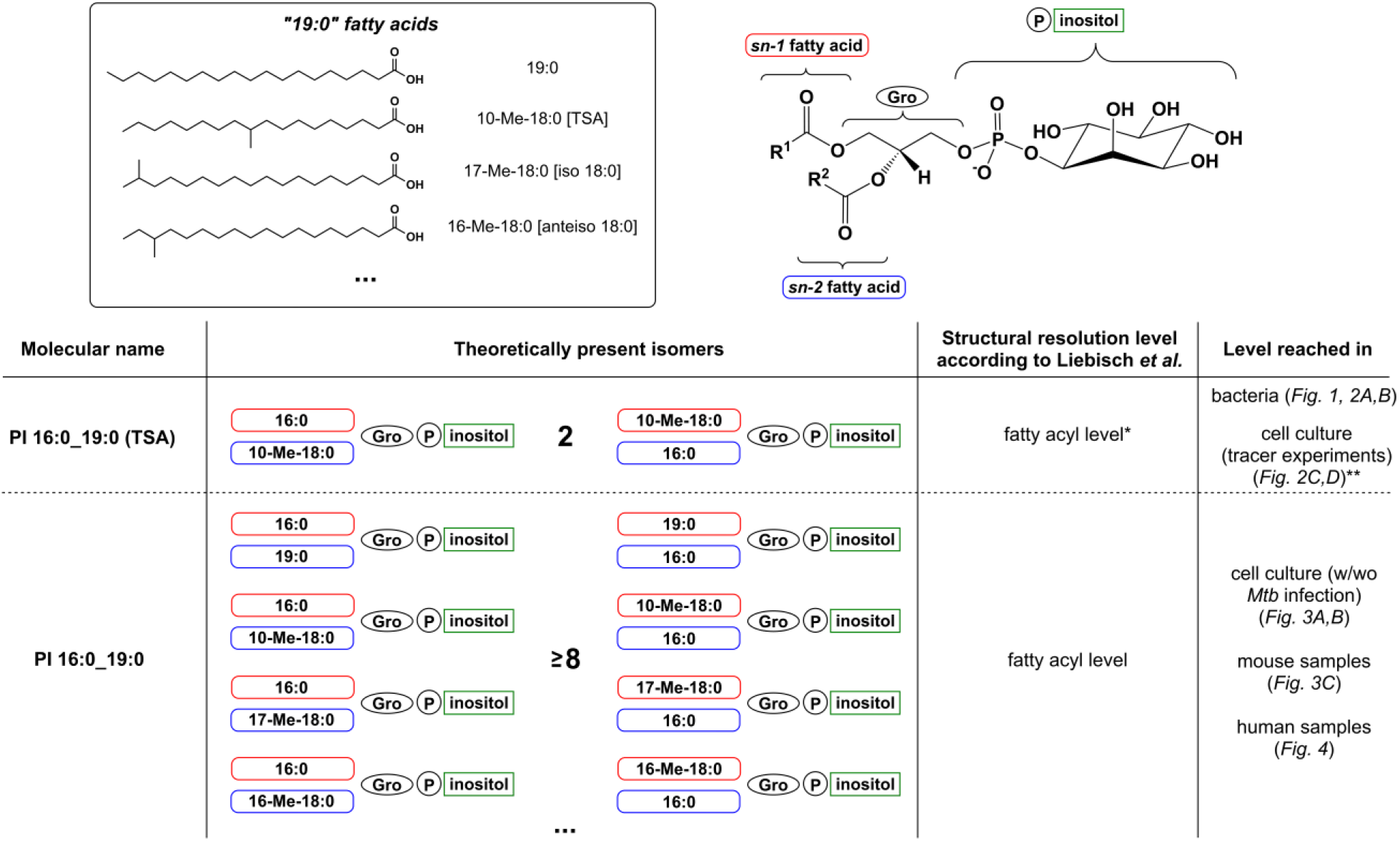
Structural complexity and analytical accuracy of the targeted MS/MS assay for PI 16:0_19:0 (TSA) in context of different biological matrices and model. * PI 16:0_19:0 (TSA) one cannot exclude *sw*-position isomerism and determine isomeric purity. However, because of the utilization of ^13^C18-labeled OA and the methylation performed by *Mtb,* we are certain of the detection of TSA. ** cell culture experiments with metabolic tracing during *Mtb* infection were demonstrated in Brandenburg et al. (*16*).

**Fig. S5.**
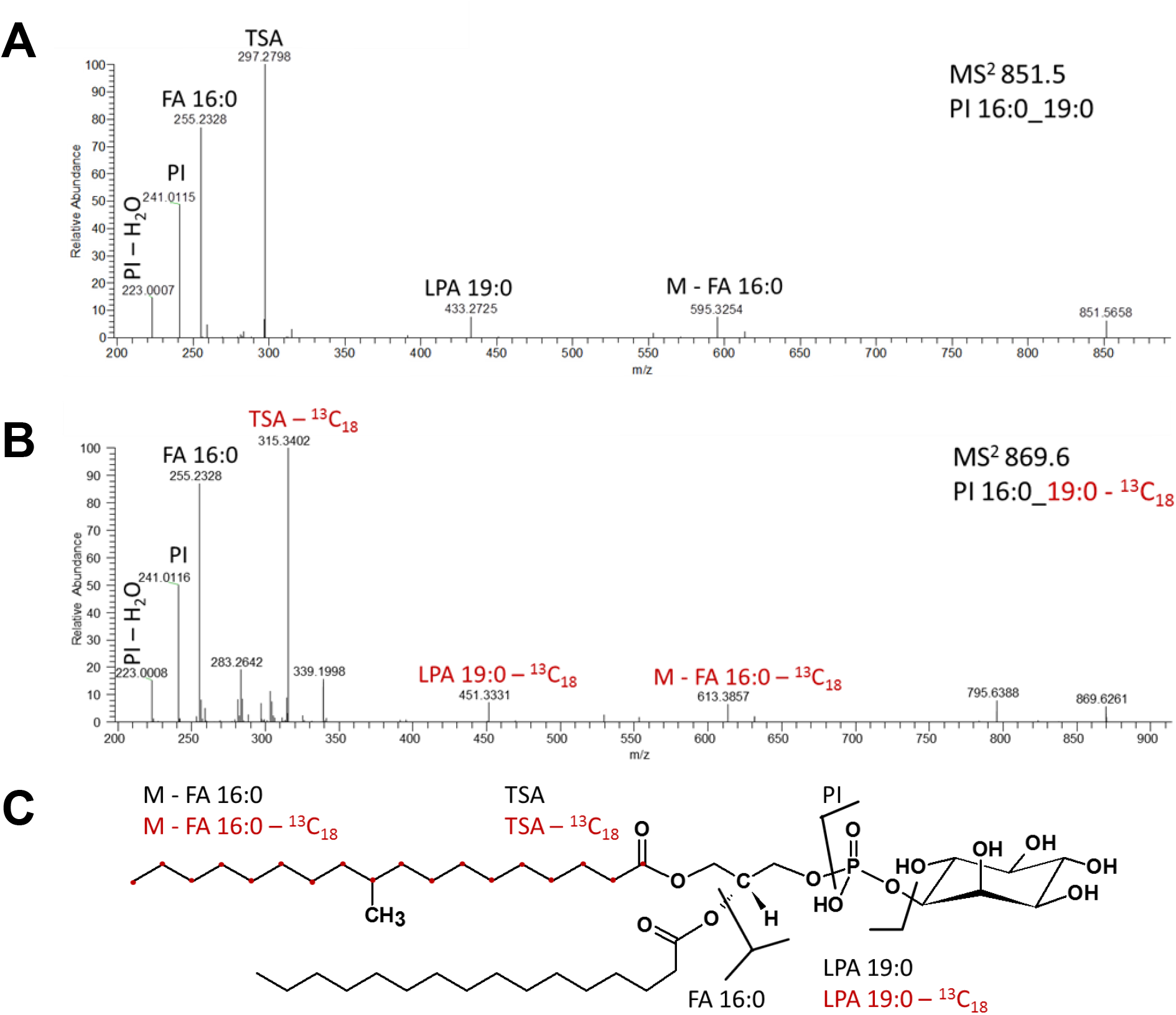
Chemical transformation of ^13^C labelled oleic acid to tuberculostearic acid (TSA) by the metabolic system of *Mtb* and structural validation of PI 16:0_19:0 (TSA) by tandem mass spectrometric analysis (MS^2^). MS^2^ spectrum of PI 16:0_19:0 (TSA) without label at precursor *m/z* 851.5 and with incorporated label. **B)** MS^2^ spectrum of *m/z* 869.6 for PI 16:0_19:0 (^13^C18-TSA). **C)** Assignment of fragment ions to chemical substructures. Fragment ions that show a shift of 18 Da due to incorporation of the ^13^C18 aliphatic chain are coloured in red.

**Fig. S6.**
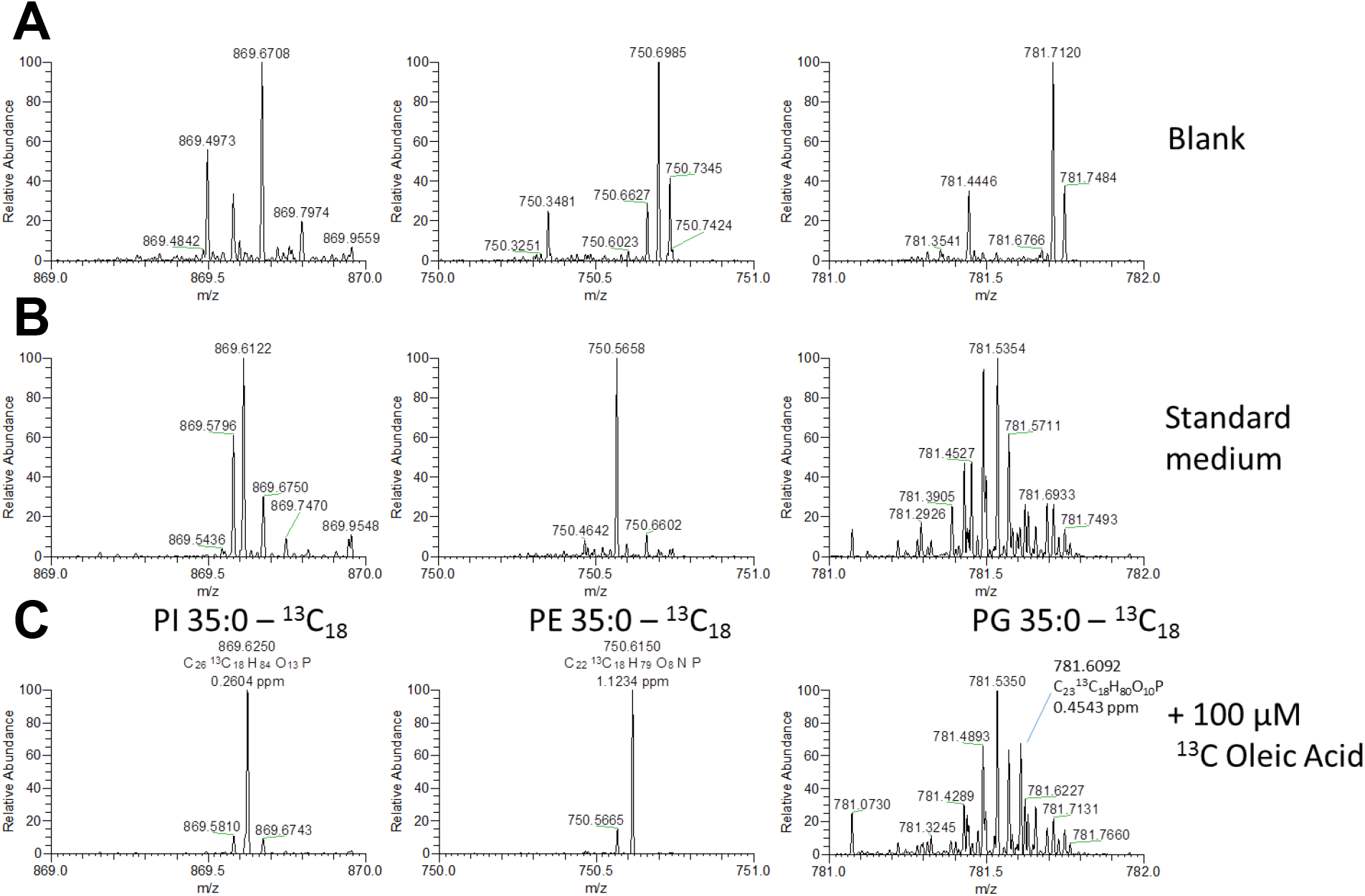
Mass spectrometric analysis of metabolic labelling of tuberculostearic acid (TSA). Representative MS^1^ *m/z* ranges for detection of labelled PI 35:0, PG 35:0 and PE 35:0 using single ion monitoring mode (SIM) for **A)** control extraction (Blank), **B)** *Mtb* grown under standard culture conditions and **C)** *Mtb* grown in media supplemented with 100 μM ^13^C labelled Oleic Acid.

**Fig. S7.**
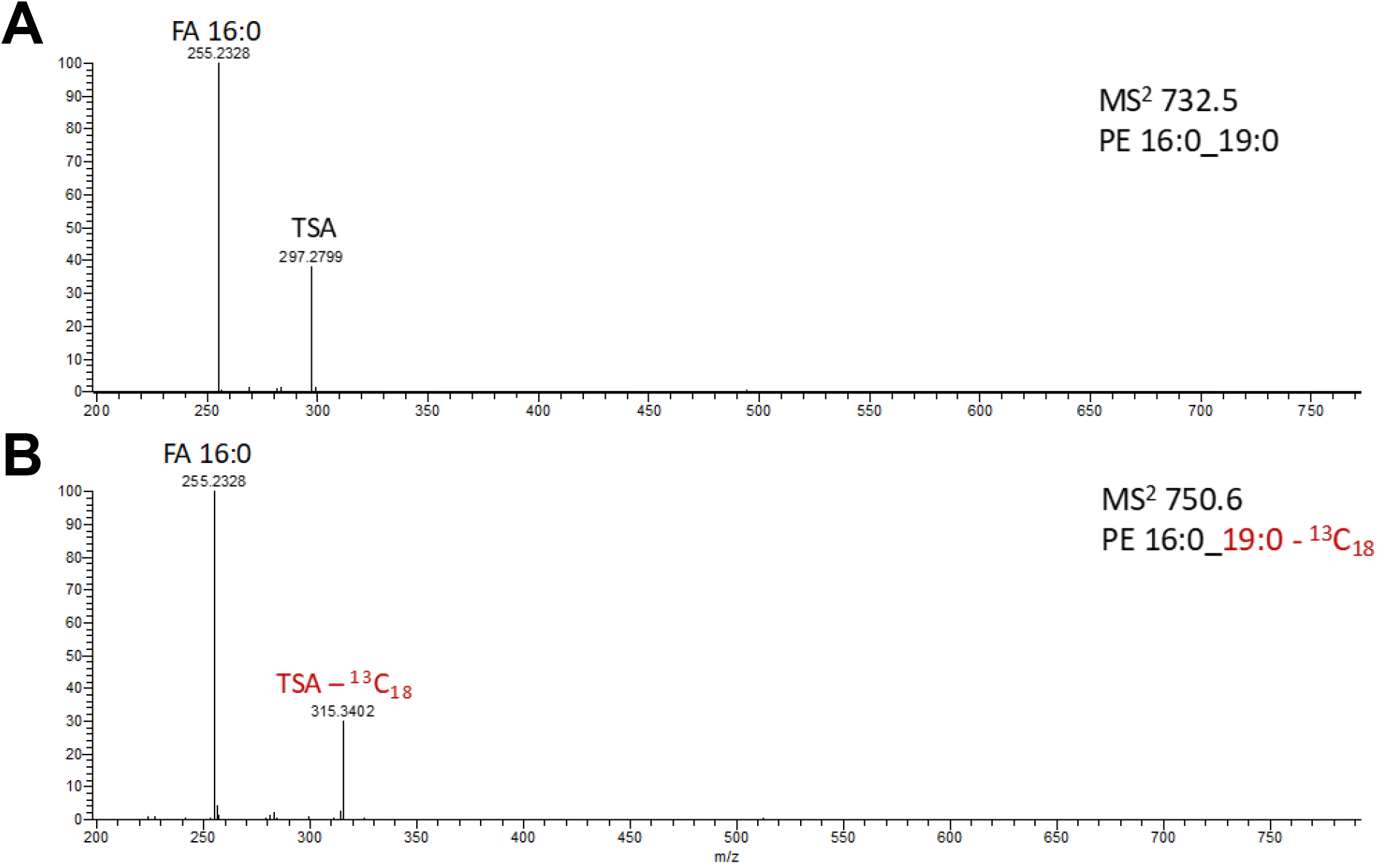
Incorporation of ^13^C18 label of oleic acid into PE 16:0_19:0 (TSA). Tandem mass spectrometric analyses (MS^2^) without ^13^C labelling at the precursor *m/z* 732.5 **A)** and with labelled Oleic Acid at *m/z* 750.6 **B)** Only the TSA fragment ion showed a shift of 18 Da due to incorporation of the ^13^C18 aliphatic chain.

**Fig. S8.**
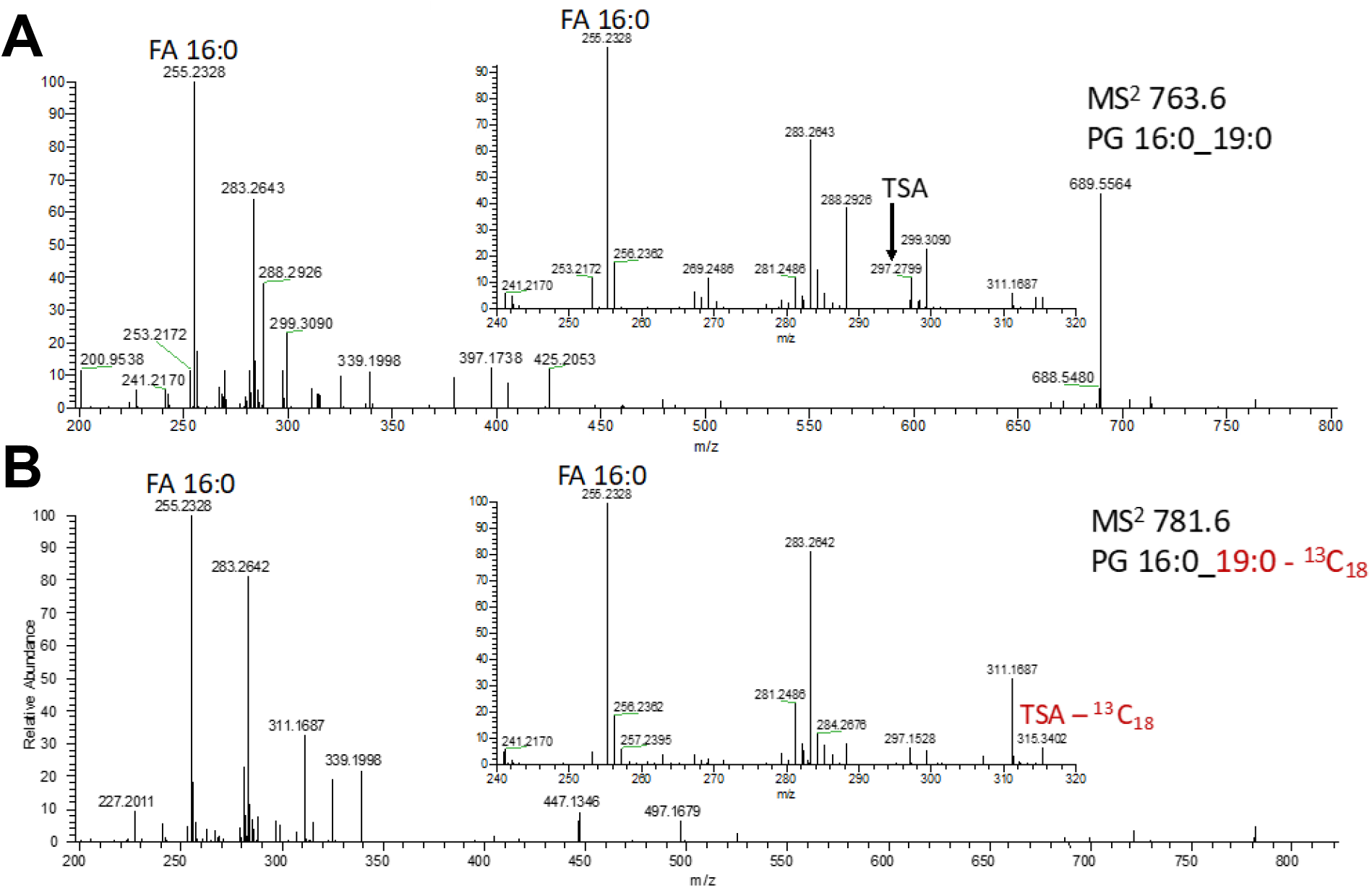
Incorporation of ^13^C18 label of Oleic Acid into PG 16:0_19:0 (TSA) by *Mtb.* Tandem mass spectrometric analyses (MS^2^) of PG 16:0_19:0 in lipid extracts of *Mtb* cultured in **A)** standard medium with the precursor *m/z* 763.6 and **B)** in media supplemented with ^13^C18 labelled Oleic Acid at *m/z* 781.6. PG 16:0_19:0 is only a minor compound for all the tested MTBC isolates. Only for the TSA fragment incorporation of the label was confirmed.

**Fig. S9.**
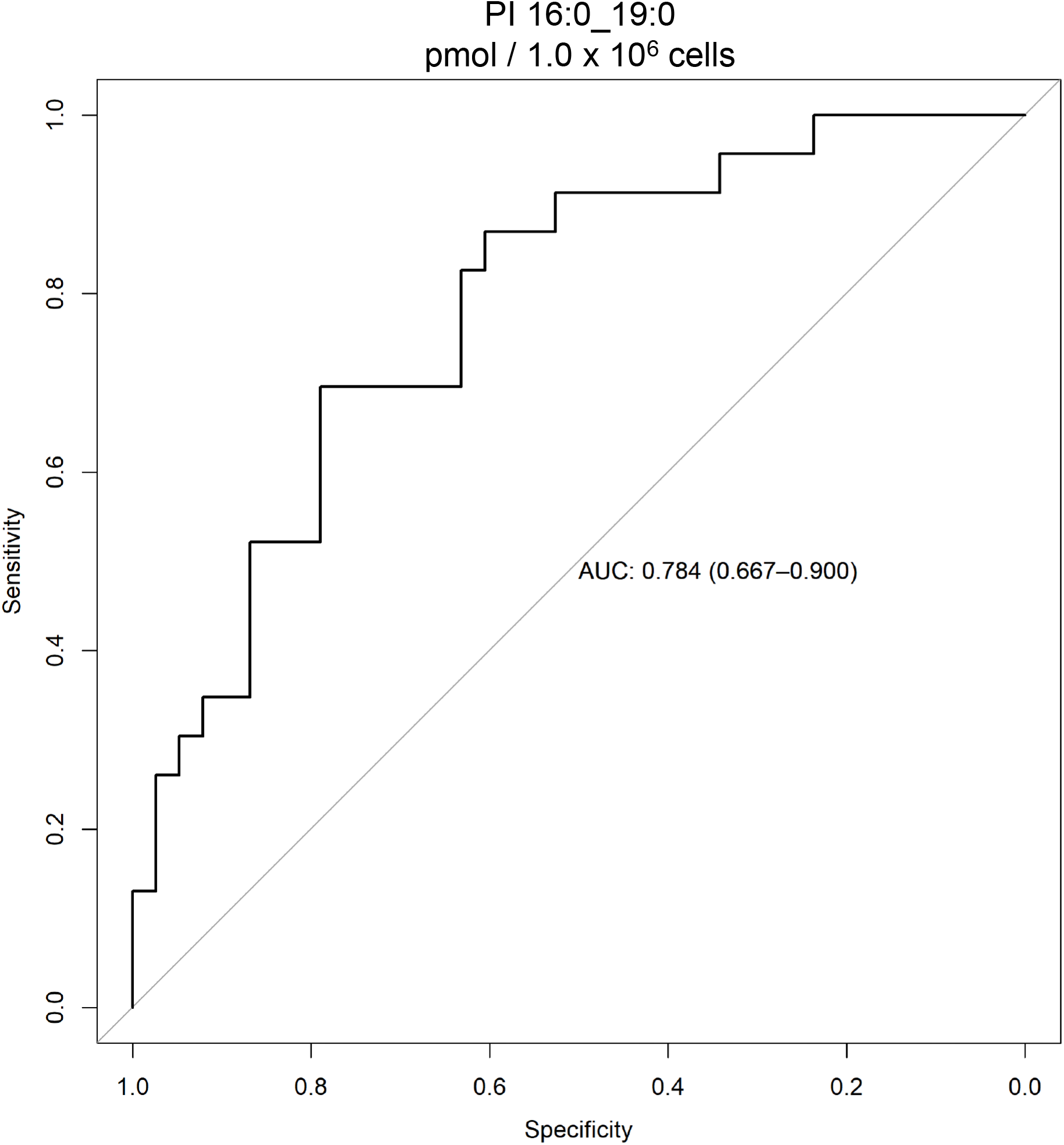
Receiver operating characteristic curve for PI 16:0_19:0 for the comparison of patients at study inclusion (t0 / tU) with healthy control. Details of the test are summarized in table S2.

**Fig. S10.**
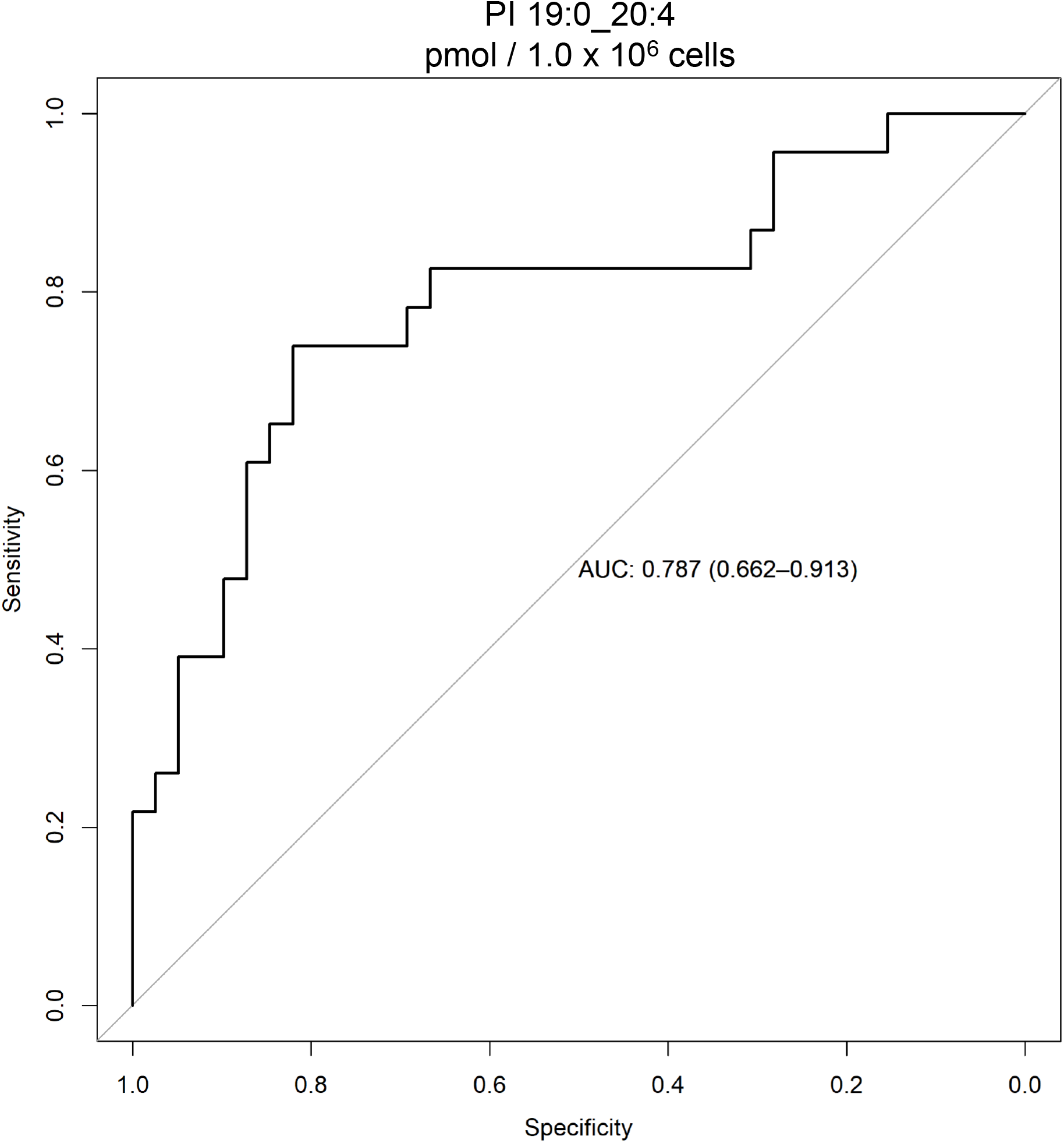
Receiver operating characteristic curve for PI 19:0_20:4 for the comparison of patients at study inclusion (t0 / tU) with healthy control. Details of the test are summarized in table S2.

**Fig. S11.**
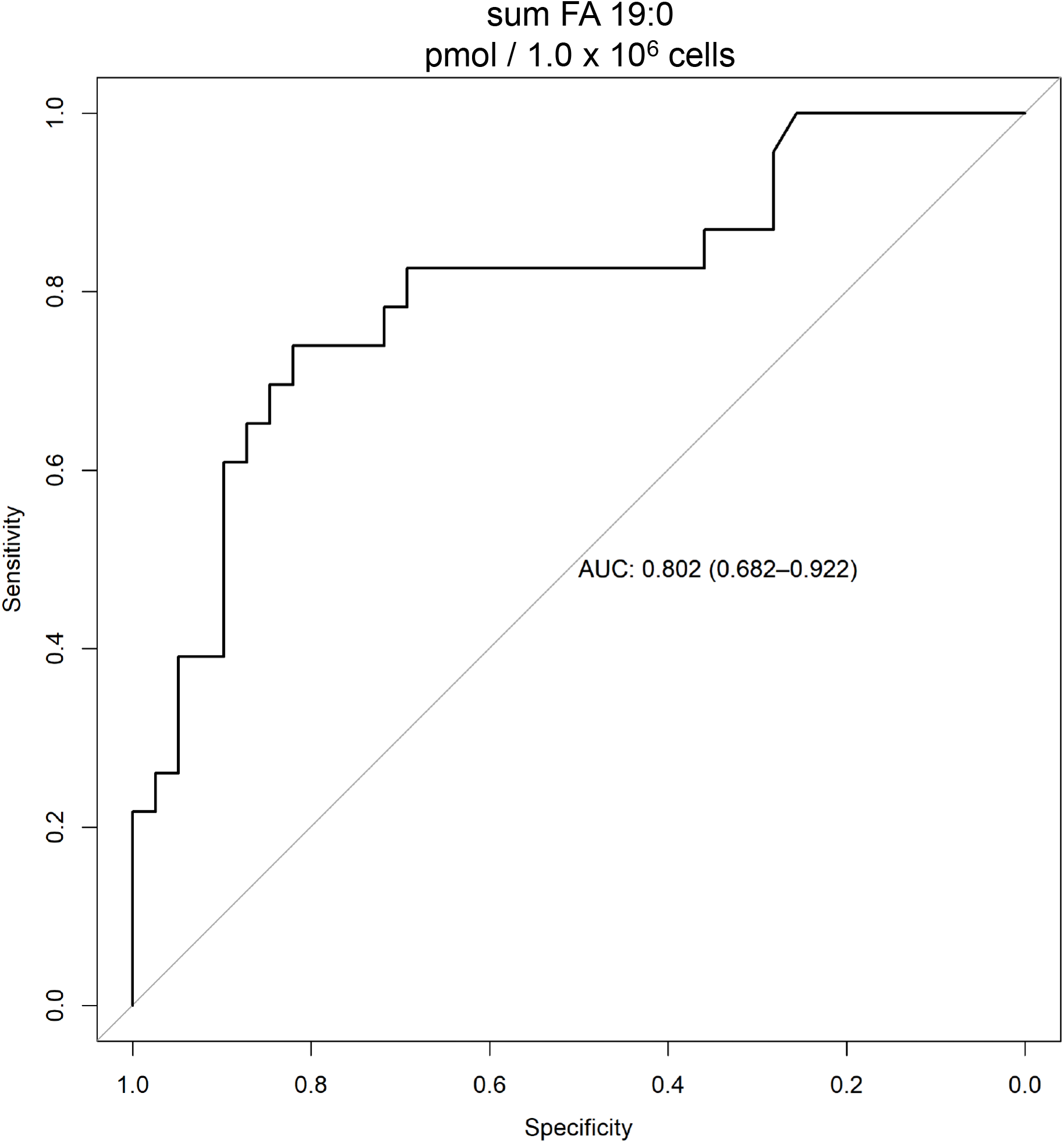
Receiver operating characteristic curve for the PI panel containing FA 19:0 (Table S3) for the comparison of patients at study inclusion (t0 / tU) with healthy control. Details of the test are summarized in table S2.

**Fig. S12.**
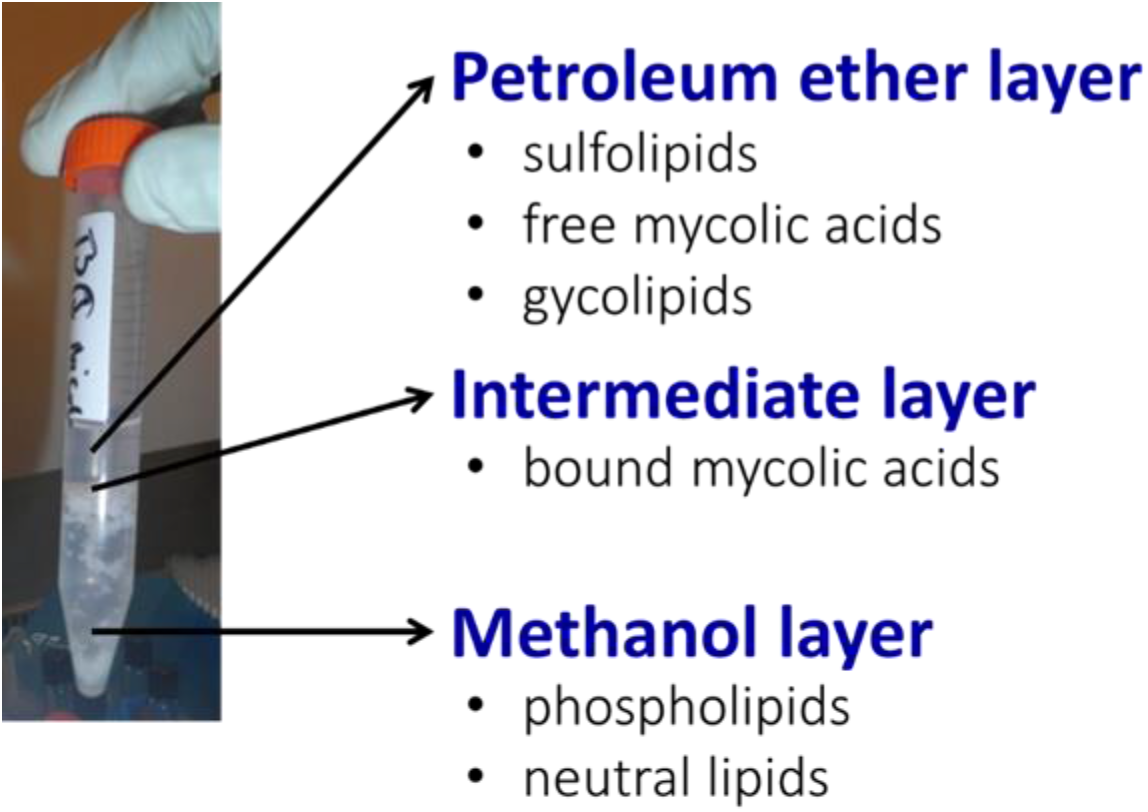
Phase separation and enriched lipid classes in the petroleum ether / methanol extraction system for MTBC strains.

**Table S1:** ROC analysis of mycobacterial derived lipid markers for TB patients at baseline and healthy control.

| Lipid marker / panel | AUC | Accuracy | Optimal Cut-off | Sensitivity | Specificity | False positive rate | False negative rate |
| --- | --- | --- | --- | --- | --- | --- | --- |
| PI 19:0_16:0 (pmol / 1.0 E06 cells) | 0.784 (0.667 – 0.900) | 0.754 | 0.228 | 0.696 | 0.790 | 0.21 | 0.30 |
| PI 19:0_20:4 pmol / 1.0 E06 cells | 0.787 (0.662 – 0.913) | 0.79 | 8.01 | 0.739 | 0.821 | 0.18 | 0.26 |
| sum FA 19:0 pmol / 1.0 E06 cells* | 0.802 (0.682 – 0.922) | 0.79 | 8.82 | 0.739 | 0.821 | 0.18 | 0.26 |

\* Panel of FA 19:0 (TSA) containing PIs listed in table S2.

**Table S2.** TSA (FA 19:0) containing PIs identified in human PBMCs utilized as marker panel.

| **exp.**<br>***m/z***<br>**(MS<sup>1</sup>)** | **Elemental**<br>**composition** | **Error in**<br>**ppm for**<br>**precursor**<br>**ion (MS<sup>1</sup>)** | **Error in ppm**<br>**for FA identification**<br>**(MS<sup>2</sup>)** | **Lipid species** | **Comment** |
| --- | --- | --- | --- | --- | --- |
| 828.5693 | *H71 C42 D7 O13*<br>*P1* | 0.0 | -0.3 and -1.8 | *PI 18:1-D7 _ 15:0* | Internal<br>standard |
| 837.5485 | H82 C43 O13 P1 | -1.6 | -0.8 and -1.9 | PI 19:0 _ 15:0 |  |
| 849.5467 | H82 C44 O13 P1 | -3.7 | -0.9 and -1.5 | PI 19:0 _ 16:1 |  |
| **851.562** | **H84 C44 O13 P1** | **-4.1** | **-0.8 and -1.0** | **PI 19:0 _ 16:0** | **Major**<br>**abundant**<br>**phospholipid**<br>**in MTBC** |
| 863.5642 | H84 C45 O13 P1 | -1.5 | -0.8 and -0.4 | PI 19:0 _ 17:1 |  |
| 875.5633 | H84 C46 O13 P1 | -2.5 | -0.8 and -0.6 | PI 19:0 _ 18:2 |  |
| 877.5775 | H86 C46 O13 P1 | -4.1 | -0.8 and -0.5 | PI 19:0 _ 18:1 |  |
| 897.551 | H82 C48 O13 P1 | 1.3 | -0.6 and -0.5 | PI 19:0 _ 20:5 |  |
| **899.5662** | **H84 C48 O13 P1** | **0.8** | **-0.7 and -0.4** | **PI 19:0 _ 20:4** | **Major**<br>**abundant**<br>**lipid of**<br>**panel** |
| 923.5577 | H84 C50 O13 P1 | -8.4 | -0.8 and -0.2 | PI 19:0 _ 22:6 |  |
| 925.5739 | H86 C50 O13 P1 | -7.9 | -0.7 and -0.1 | PI 19:0 _ 22:5 |  |

**Table S3.** Components of the internal standard mix for lipid analysis of human PBMCs.

| **Internal standard components** | **pmol/sample (10 μL)** | **pmol/sample (20 μL)** |
| --- | --- | --- |
| 15:0-18:1(d7) PC | 83.65 | 167.3 |
| 15:0-18:1(d7) PE | 3.14 | 6.28 |
| 15:0-18:1(d7) PS (Na-Salt) | 2.12 | 4.24 |
| 15:0-18:1(d7) PG(Na-Salt) | 14.94 | 29.88 |
| 15:0-18:1(d7) PI(NH<sub>4</sub>-Salt) | 4.19 | 8.38 |
| 15:0-18:1(d7) PA(Na-Salt) | 4.19 | 8.38 |
| 18:1(d7) LPC | 18.89 | 37.78 |
| 18:1(d7) LPE | 4.27 | 8.54 |
| 18:1(d7) Chol Ester | 212.11 | 424.22 |
| 18:1(d7) MG | 2.16 | 4.32 |
| 15:0-18:1(d7) DG | 6.27 | 12.54 |
| 15:0-18:1(d7)-18:1 TG | 27.64 | 55.28 |
| 18:1(d9) SM | 16.43 | 32.86 |
| Cholesterol(d7) | 97.96 | 195.92 |
| Ceramide C17 | 37.68 | 75.36 |

Internal standard mix is composed of C17 ceramide and SPLASH mixture purchase from Avanti Polar Lipids (Alabaster, US). For samples with cell counts of $1 \times 10^6$ and $2 \times 10^6$ 10 μL of the internal standard were added prior extraction. For samples containing the maximal cell counts of $5 \times 10^6$ a volume of 20 μL of the standard mix were utilized.

**Table S4:** Gradient and flow rates used for the separation of the lipids by LC-MS.

| **Time (min)** | **% A** | **% B** | **Flow (μL / min)** |
| --- | --- | --- | --- |
| 0 | 0 | 100 | 15 |
| 1.5 | 0 | 100 | 15 |
| 8.2 | 70 | 30 | 15 |
| 11.66 | 70 | 30 | 15 |
| 12 | 80 | 20 | 15 |
| 13.3 | 80 | 20 | 15 |
| 13.66 | 0 | 100 | 15 |

Solvents: **A**, chloroform, methanol and ammonia solution 28-30% (86/13/1, v/v/v); **B**, chloroform, methanol and ammonia solution 28-30% (50/49/1, v/v/v)

Data file S1. Major abundant extractable membrane lipids of MTBC strains.

Data file S2. Details of targeted lipid analysis in mice infection model including standardization of lung tissue homogenate extraction procedure.

Data file S3. Clinical data and lipidomics results of healthy study participants and TB patients.

Data file S4. Clinical data and lipidomics results of all TB patients included in this study at therapy start (t0).

Data file S5. Clinical data and lipidomics results of all TB patients grouped according to the WHO and TBnet outcome criteria at therapy start (t0) and end (tE).

